# Homeobox Transcription Factor HbxA Influences Expression of over One Thousand Genes in the Model Fungus *Aspergillus nidulans*

**DOI:** 10.1101/2023.03.30.533655

**Authors:** S.S. Pandit, J. Zheng, Y. Yi, S. Lorber, O. Puel, S. Dhingra, E.A. Espeso, A.M Calvo

## Abstract

In fungi, conserved homeobox-domain (HD) proteins are transcriptional regulators governing development. In *Aspergillus* species, several HD transcription factor genes have been identified, among them, *hbxA*/*hbx1*. For instance, in the opportunistic human pathogen *Aspergillus fumigatus*, *hbxA* is involved in conidial production and germination, as well as virulence and secondary metabolism (SM), including production of fumigaclavines, fumiquinazolines, and chaetominine. In the agriculturally important fungus *Aspergillus flavus,* disruption of *hbx1* results in fluffy aconidial colonies unable to produce sclerotia. *hbx1* also regulates production of aflatoxins, cyclopiazonic acid and aflatrem. Furthermore, transcriptome studies revealed that *hbx1* has a broad effect on the *A. flavus* genome, including numerous genes involved in SM. These studies underline the importance of the HbxA/Hbx1 regulator, not only in developmental processes but also in the biosynthesis of a broad number of fungal natural products, including potential medical drugs and mycotoxins. To gain further insight into the regulatory scope of HbxA in *Aspergilli*, we studied its role in the model fungus *Aspergillus nidulans*. Our present study of the *A. nidulans hbxA*-dependent transcriptome revealed that more than one thousand genes are differentially expressed when this regulator was not transcribed at wild-type levels, among them numerous transcription factors, including those involved in development as well as in SM regulation. Furthermore, our metabolomics analyses revealed that production of several secondary metabolites, some of them associated with *A. nidulans hbxA*-dependent gene clusters, was also altered in deletion and overexpression *hbxA* strains compared to the wild type, including synthesis of nidulanins A, B and D, versicolorin A, sterigmatocystin, austinol, dehydroaustinol, and three unknown novel compounds.

## INTRODUCTION

Developmental studies of the model filamentous fungus *Aspergillus nidulans* have provided broad valuable insight into the genetic regulatory mechanisms of morphogenesis in fungi (1–3). *A. nidulans* efficiently disseminates by asexual reproduction, forming specialized structures called conidiophores, which bares large numbers of air-borne conidia. Activation of conidiogenesis is mediated by several transcription factor genes, including *flb* genes, such as *flbB*, *flbC*, *flbD*, and *flbE*, which activate the central regulatory pathway comprised of *brlA*, *abaA* and *wetA* (4). This model organism is also able to reproduce sexually by producing cleistothecia, fruiting bodies containing meiospores called ascospores. Cleistothecia form by aggregation of vegetative mycelia, surrounded by nursing Hülle cells. This results in the formation of cleistothecial primordia, which later mature into melanized cleistothecia (5,6). Several genes are involved in the regulation of these processes, including *nsdD*, *medA*, *phoA*, *stuA*, *lsdA* and *tubB* (7–12).

Other developmental regulators include Homeobox-domain transcription factors (HD-TFs). These are global regulators governing developmental processes in many eukaryotic organisms (13–15). The HD contains approximately 66 conserved amino acid that bind to the promoter of genes governing development and other cellular processes in fungi, plants, and animals. In general, fungi possess 6–12 HD-TF genes in their genome (13,14,16,17). The first reported HD-FT gene is *pah1* in *Podospora anserina*, where it controls microconidiation as well as mycelial branching (18). Loss-of-function of seven HD-TF genes in this fungus revealed their role in sexual development (19). Another study showed that several HD-TFs in the rice pathogen *Magnaporthe oryzae* are necessary for proper hyphal growth, asexual development, and appressorium formation (20,21). In three species of *Fusarium*, loss-of-function of the *htf1* homeobox gene leads to alteration of phialides during conidiophore formation, accompanied by a drastic reduction in conidial production (22). In the fungus *Botrytis cinerea,* the BcHOX8 gene has been shown to regulate growth, conidiation, and virulence in different host plants (16). Also, lack of the *GRF10* HD-TF gene in the human pathogen *Candida albicans* resulted in a decrease in growth, defects in chlamydospore morphology, alterations in biofilm production, and a reduction of virulence (23).

HD-TFs are also key regulators in species of the genus *Aspergillus*. In the agriculturally relevant fungus *Aspergillus flavus*, deletion of eight HD-FT genes revealed that *hbx1* in particular, was required for normal vegetative growth and production of conidia and sclerotia. The regulation of morphological development as well as regulation of SM, are often genetically linked (24–26). Interestingly, in this case also, the production of secondary metabolites, including mycotoxins (aflatoxins, cyclopiazonic acid and aflatrem), was under the regulation of *hbx1* (17). Furthermore, study of the *hbx1-*dependent transcriptome indicated its importance in morphological development and in regulation of secondary metabolite production (27). Remarkably, the gene category corresponding to SM was the most affected by *hbx1*. Additionally, in our previous study of the *hbx1* homolog in the opportunistic human pathogen *Aspergillus fumigatus*, *hbxA*, showed that this gene is necessary for proper spore formation, regulating expression of *brlA, flbB, flbD* and *fluG* (28). The *hbxA* gene also influenced germination rate and virulence in a neutropenic mouse model. Interestingly, as in the case of *A. flavus*, *A. fumigatus hbxA* affected production of various secondary metabolites, including fumigaclavines, fumiquinazolines, compounds that accumulate in asexual structures, whose production is linked to *brlA* expression (29–32), and chaetominine, an alkaloid compound that is being tested to combat leukemia cells(20). Both *A. flavus* and *A. fumigatus* studies indicate that HbxA/Hbx1 is a global regulator of SM in these fungi, in addition to its role in morphogenesis. HbxA also affects *A. nidulans* conidiation (33,34) in a similar manner as that in *A. flavus* and *A. fumigatus*(17,27,28). To gain further inside into the regulatory scope of *hbxA* in the genus *Aspergillus*, in the present study, we characterized its role in the model fungus *A. nidulans* by transcriptome and metabolomics approaches. Our findings indicate that more than one thousand genes were differentially expressed in the absence of this regulator or when it was over-expressed, as compared to the wild type. These include several transcription factor genes, including those involved in development and SM production. Our study revealed that numerous secondary metabolites gene clusters are *hbxA*-dependent in *A. nidulans*. Furthermore, our analyses also indicated that *A. nidulans* metabolome is affected by *hbxA*, including production of some unknown novel compounds.

## MATERIALS and METHODS

### Phylogenetic Analysis

Deduced amino acid sequences of HbxA homologs were obtained from FUNGIDB (https://fungidb.org/fungidb/) website. BLASTp was performed against the protein sequence database (pdb). Percentage (%) similarity was found using Pairwise sequence alignment using EMBOSS Needle (ebi.ac.uk/Tools/psa/emboss_needle/). The phylogenetic tree was constructed using MEGA v6.0 and the Maximum Likelihood model with bootstrap value of 1000.

### Strains used and culture conditions

The *A. nidulans* strains used in this study are listed in Table 1. Strains were grown on glucose minimal medium (GMM) (35) with appropriate supplements for their respective auxotrophic markers (35). For solid medium, agar (15 g/L) was added. Strains were stored as 30% glycerol stocks at −80°C.

**Table 1:** Strains used in this study

| Strain Name | Pertinent Genotype | Source |
| --- | --- | --- |
| RMJP1.49 | *pyrG89; argB2; Δnku::argB; pyroA4* | (36) |
| TSSP38.1 | *pyrG89; argB2; Δnku::argB; pyroA4;ΔhbxA::pyrG;pyroA* | This Study |
| TSSP40.1 | *pyrG89; argB2; ΔnkuA::argB; pyroA4; ΔhbxA::pyrG, hbxA::pyroA* | This Study |
| TSSP34.1 | *pyrG89; argB2; ΔnkuA::argB; pyroA4; gpdA(p)::ΔhbxA::trpC(t)::pyrG; pyroA* | This Study |
| TRV50.2 | Wild type | (37) |

### Generation of the hbxA deletion strain (ΔhbxA)

The DNA cassette employed to obtain the deletion *hbxA* strain (TSSP38.1) was generated by fusion polymerase chain reaction (PCR) through a previously described method (38). All primers used in this study are listed in Table 2. The 1.5 kb 5’ UTR region of the *hbxA* locus was PCR amplified using P#2154/SD3 and P#2155 primers from genomic DNA of the *A. nidulans* FGSC4 wild-type strain. Similarly, the 1.1 kb 3’ UTR of *hbxA* was amplified using P#2156 and P#2157 primers also from genomic DNA. The 1.9 kb *A. fumigatus pyrG* selectable marker was amplified from plasmid p1439 (39) using P#2158 and P#2159 primers. The 5’ and 3’ UTR fragments were fused to the selectable *pyrG* marker using P#2160/SD9 and P#2161 primers. The resultant fusion product was transformed into RMJP1.49 strain using a polyethylene glycol mediated protocol as described previously (38). Transformants were confirmed by diagnostic PCR using P#2154 and P#963 primers. The selected deletion *hbxA* strain was then transformed with a DNA fragment containing the *A. nidulans pyroA* gene, PCR amplified with primers P#1042 and P#1045 from genomic DNA, resulting in strain TSSP38.1.

**Table 2:**
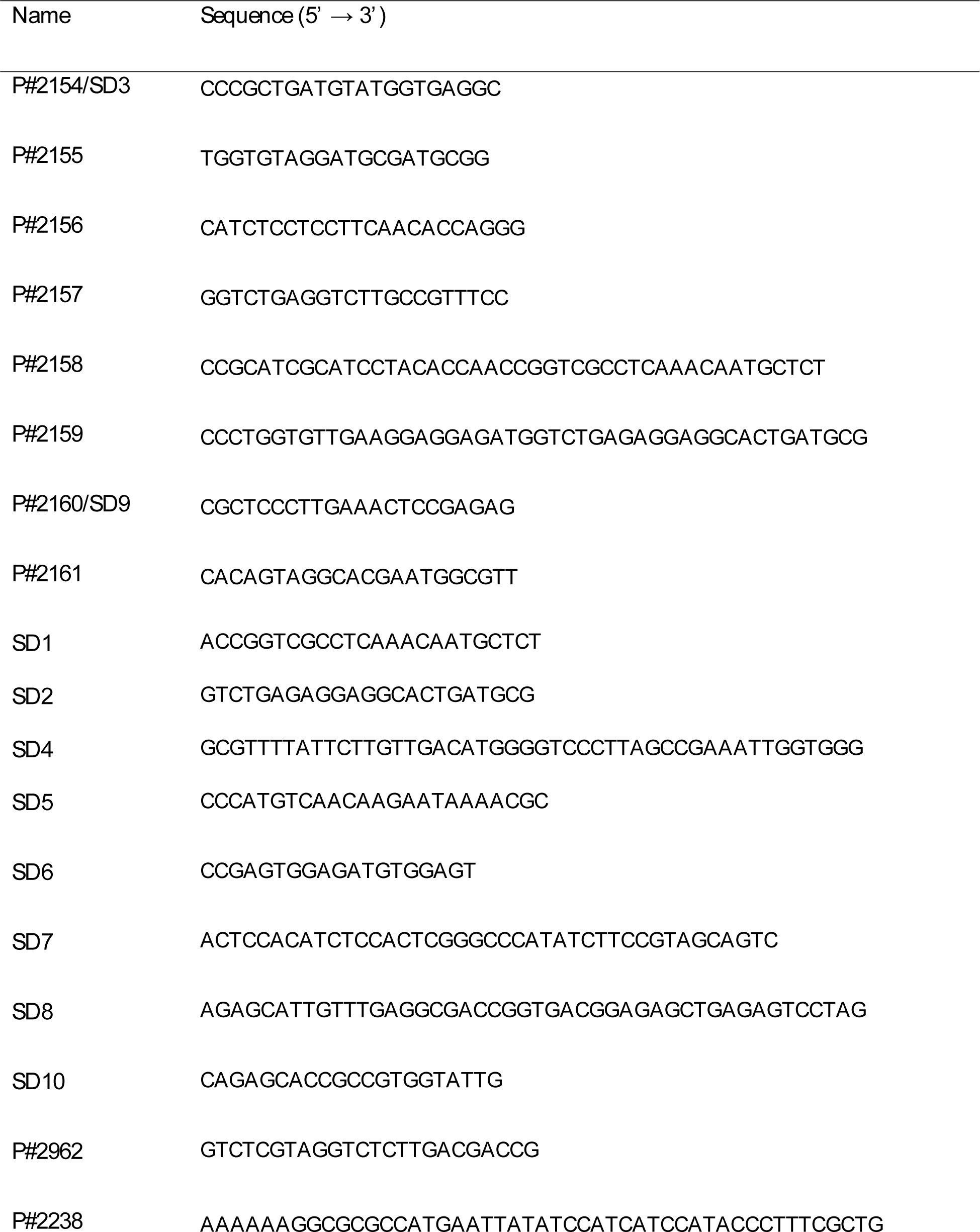
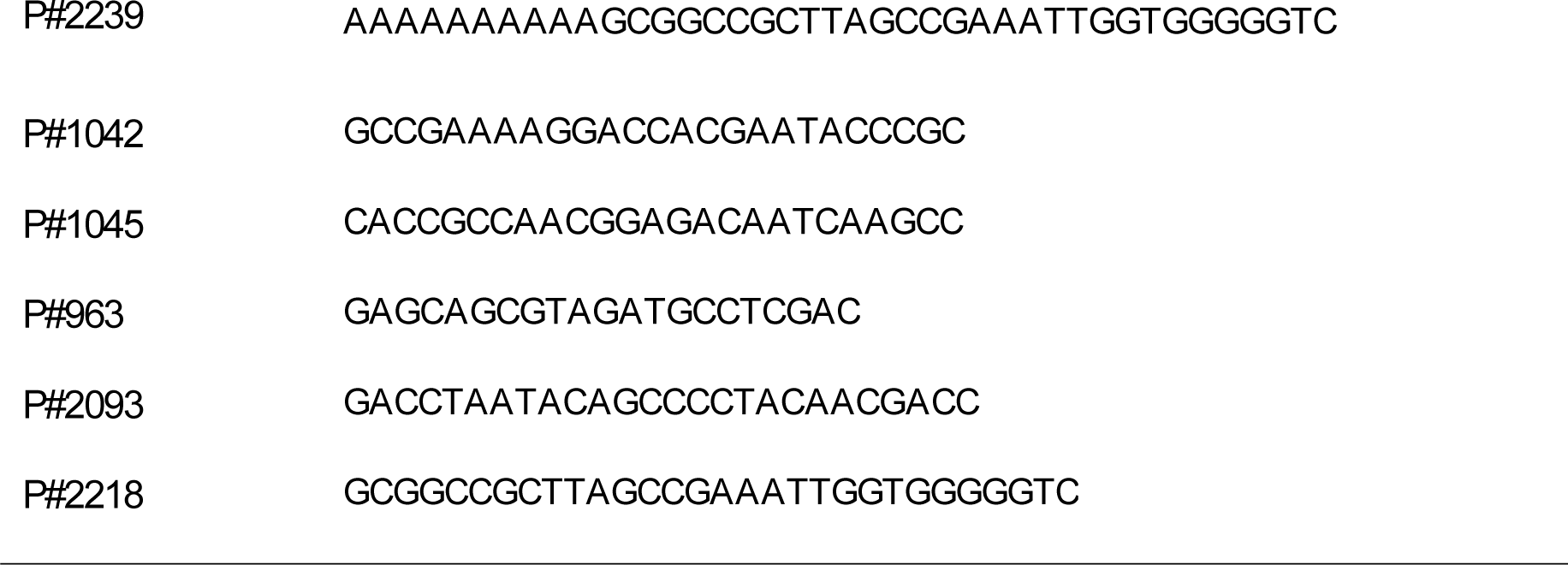
Primers used in this study

### Generation of the hbxA complementation strain (hbxA-com)

The complementation strain (TSSP40.1) was generated by re-introducing the wild-type *hbxA* allele into the Δ*hbxA* strain at the same locus. The complementation cassette was generated as follows: first, a DNA fragment containing the *hbxA* coding region and a 3.7 kb 5’UTR was PCR amplified using P#2154/SD3 and SD4, and the *trpC* terminator fragment was amplified with primers SD5 and SD6 using *A. nidulans* genomic DNA as a template. The *A. fumigatus pyroA* gene (Afub_055620) was amplified from genomic DNA using primers SD7 and SD8. *A. fumigatus pyrG* was amplified from plasmid p1439 (38) using primers SD1 and SD2. All four PCR fragments were fused together using primers P#2160/SD9 and SD10 in a single reaction using Prime Star DNA polymerase (Clonetech, USA). The resulting fusion product was then transformed into the *hbxA* deletion strain (TSSP38.1) using methods previously described (38). Fungal transformants were confirmed using diagnostic PCR with primers P#2154 and P#2962.

### Generation of the hbxA overexpression strain *(OE*hbxA)

To generate the over-expression *hbx1* strain (TSSP34.1), the coding region of *hbxA* was first amplified from *A. nidulans* genomic DNA using P#2238 and P#2239 primers. The resulting PCR product was digested with *Asc*I and *Not*I and ligated to pTRS2 plasmid, previously digested with the same enzymes. pTRS2 contains the *gpdA* promoter, *gpdA*_(p)_, and *trp*C terminator, *trp*C_(t)_. The resulting plasmid, pSSP34.1, was transformed into the *A. nidulans* RJMP1.49 strain, and transformants were screened by PCR using P#2093 and P#2218 primers. The selected overexpression *hbxA* strain was then transformed with a DNA fragment containing the *A. nidulans pyroA* gene, PCR amplified with primers P#1042 and P#1045 from genomic DNA, resulting in strain TSSP34.1.

### Transcriptome analysis

#### RNA purification and sequencing

Plates containing 25 mL of solid GMM with the appropriate supplements were top-agar inoculated with 5 mL of medium containing ∼5 x10^6^ spores/mL of wild-type (WT) control, Δ*hbxA*, *hbxA-*com or OE*hbxA* (Table 1). Cultures were incubated in the dark at 37°C. After 72 h of incubation, mycelia were collected, frozen in liquid nitrogen, and lyophilized. Total RNA was extracted from mycelia using an RNeasy Plant Mini Kit (Qiagen, Germantown, Maryland, USA) following the manufacturer’s protocol. RNA was further purified using Dynabeads mRNA Purification Kit (Thermo Fisher Scientific Inc., Massachusetts, USA). RNA quality was assessed using an Agilent Bioanalyzer. Sequencing was performed as a HiSeq 2000 single read 1×100bp lane. The experiment was carried out with 3 biological replicates.

#### Read mapping, decontamination and Read count

The RNA reads were trimmed by trim_galore (40) with the default parameter. Kraken2 (41) was run on trimmed reads to check the contamination. Then, reads were mapped to reference genome downloaded from FungiDB (*Aspergillus nidulans* FGSC4)(42) . Unmapped reads were removed to get clean reads. The clean reads were then repaired to pair-end reads with BBTools (43). These final clean pair-end reads were remapped to reference genome again using hisat2 ((44).

Mapped reads in SAM format were sorted by coordinates with samtools (45) to obtain the BAM format mapped reads. Then read count and TPM (Transcripts Per Kilobase Million) were calculated by running StringTie (46) and python script. The parameters were set not to infer new transcripts with the reference gene annotation file (also downloaded from FungiDB).

#### Differentially expressed coding genes (DEGs)

The read counts table was used as input for DEseq2 (47). This package was used to determine DEGs by comparing read counts between two strains. Significant up regulated genes were determined with −log10 q-value <= 2 and log2 fold change >= 2, while significant down regulated genes were defined with −log10 q-value <= 2 and log2 fold change <= −2. Control vs. OE*hbxA* and Control vs. Δ*hbxA*. python script was developed to convert gene id between FungiDB and FungiFun2 so that the webserver of FungiFun2 can be used to perform FunCat term annotation and enrichment of DEGs for Control vs. OE*hbxA* and Control vs. Δ*hbxA*(48). Heat maps of TPM (transcript per million) values of DEGs of secondary metabolism clusters were calculated by averaging all TPM values of all replicates.

Evaluation of differentially expressed ortholog genes in *A. nidulans* and *A. flavus* was carried out by using the MCL algorithm in combination with all-versus-all protein BLAST search, similar to a method previously described (49). Proteins with BLAST hits were filtered with the following parameters: 1, query and subject coverage is greater than 60%. 2, e-value is less than 1^-5^. 3, the percent of identity is greater than 60%. And then, the filtered hits were fed into OrthoMCL with an inflation parameter of 2 to generate orthogroups between these two species.

To analyze changes in the expression of genes in secondary metabolite biosynthetic gene clusters (SMGs), 67 SMGs were extracted (50). SMGs expression related figures were plotted with python seaborn package. In addition, expression of 521 transcript factors (TFs) was also analyzed. The list of TFs and their function annotations were derived from a previous report (51).

### Metabolomics

#### Thin-Layer Chromatography

Wild-type control, Δ*hbxA*, *hbxA-*com, and OE*hbxA* were top-agar inoculated with 5 mL of medium containing ∼5 x10^6^ spores/mL on solid GMM and grown at 37°C for 3 days. Three 16-mm diameter cores per plate were collected and extracted with chloroform. Overnight dried extracts were resuspended in 200 µL chloroform. Sample were separated using thin-layer chromatography (TLC) as previously described (28,52) on silica gel plates using benzene and glacial acetic acid [95:5(*v*/*v*)] as solvent system. Aluminum chloride (15% in ethanol) was then sprayed, and plates were baked for 10 min at 80 °C. Bands were visualized under UV light (375 nm). Sterigmatocystin (ST) standard was purchased from Sigma-Aldrich (St. Louis, MO, USA).

#### Analysis of secondary metabolites by liquid chromatography combined with mass spectrometry (LC-MS)

Chloroform extracted samples were also analyzed by LC-MS. Samples were resuspended in 500 μL of acetonitrile/water (50:50, v/v), shaken vigorously for 30 s and then treated with a sonicator (Bransonic 221 Ultrasonic bath, Roucaire, Les Ulis, France) for 2 h. A volume of 250 µL of pure ACN was added to each sample, followed by vigorous shaking (30s) and centrifugation (pulse). Secondary metabolites analysis was performed using Acquity ArcSystem HPLC (Waters, Saint-Quentin-en-Yvelines, France) combined with an LTQ Orbitrap XL high-resolution mass spectrometer (Thermo Fisher Scientific, Les Ulis, France). A volume of 10 μL of the suspension was injected into a reversed-phase 150 mm × 2.0 mm, Luna® 5 μm C18 column (Phenomenex, Torrance, CA, U.S.A.). Water acidified with 0.1% formic acid was used as phase A and 100% acetonitrile was used as phase B with the following elution gradient: 0 min 20% B, 30 min 50% B, from 35 to 45 min 90% B, from 50 to 60 min 20% B at 30 °C at a flow rate of 0.2 mL min^-1^. HRMS acquisitions were achieved with electrospray ionization (ESI) in positive and negative modes, as previously reported (28). MS/MS spectra were obtained with CID mode at low resolution and collision energy of 35%.

### Statistical analysis

Statistical analysis was applied to analyze all quantitative data in this study utilizing analysis of variance (ANOVA) in conjunction with a Tukey multiple-comparison test using a *p* value of <0.05 for samples that are determined to be significantly different.

## RESULTS

### HbxA is conserved in numerous fungal species

Our phylogenetic analysis confirmed that the *hbxA* deduced amino acid sequence corresponds to a transcription factor containing a homeodomain. HbxA homologs are present in other *Aspergillus* species, including *A. flavus* (17,27), *A. fumigatus* (28), *Aspergillus niger* and *Aspergillus terreus* (Fig 1, Table 3), as well as in species of other fungal genera, such as *Alternaria alternata, Arthrobotrys flagrans, Ascosphaera apis, Blastomyces dermatitidis, Histoplasma capsulatum, Microsporum canis, Penicilliopsis zonata, Penicillium rubens, Talaromyces marneffei* and *Trichophyton tonsurans* (Fig 1, Table 3). Of the sequences analyzed, *A. niger* HbxA was the closest homolog to *A. nidulans* HbxA, with 56.40% identity and 68.4% sequence similarity.

**Fig 1:**
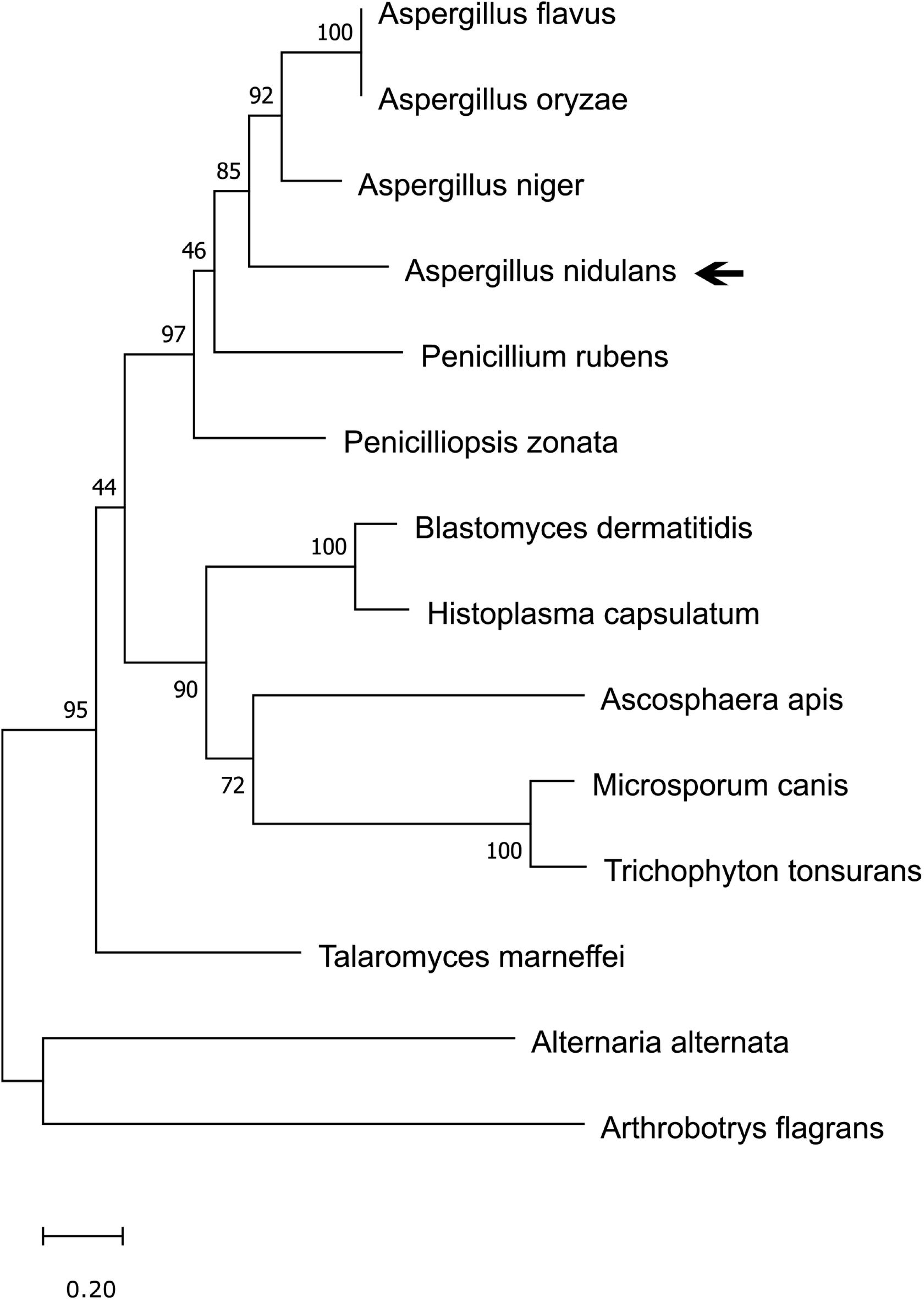
Phylogenetic analysis of *Aspergillus nidulans* HbxA. The phylogenetic tree was constructed using MEGA v6.0 and the Maximum Likelihood model with bootstrap value of 1000 (http://megasoftware.net/).

**Fig 2:**
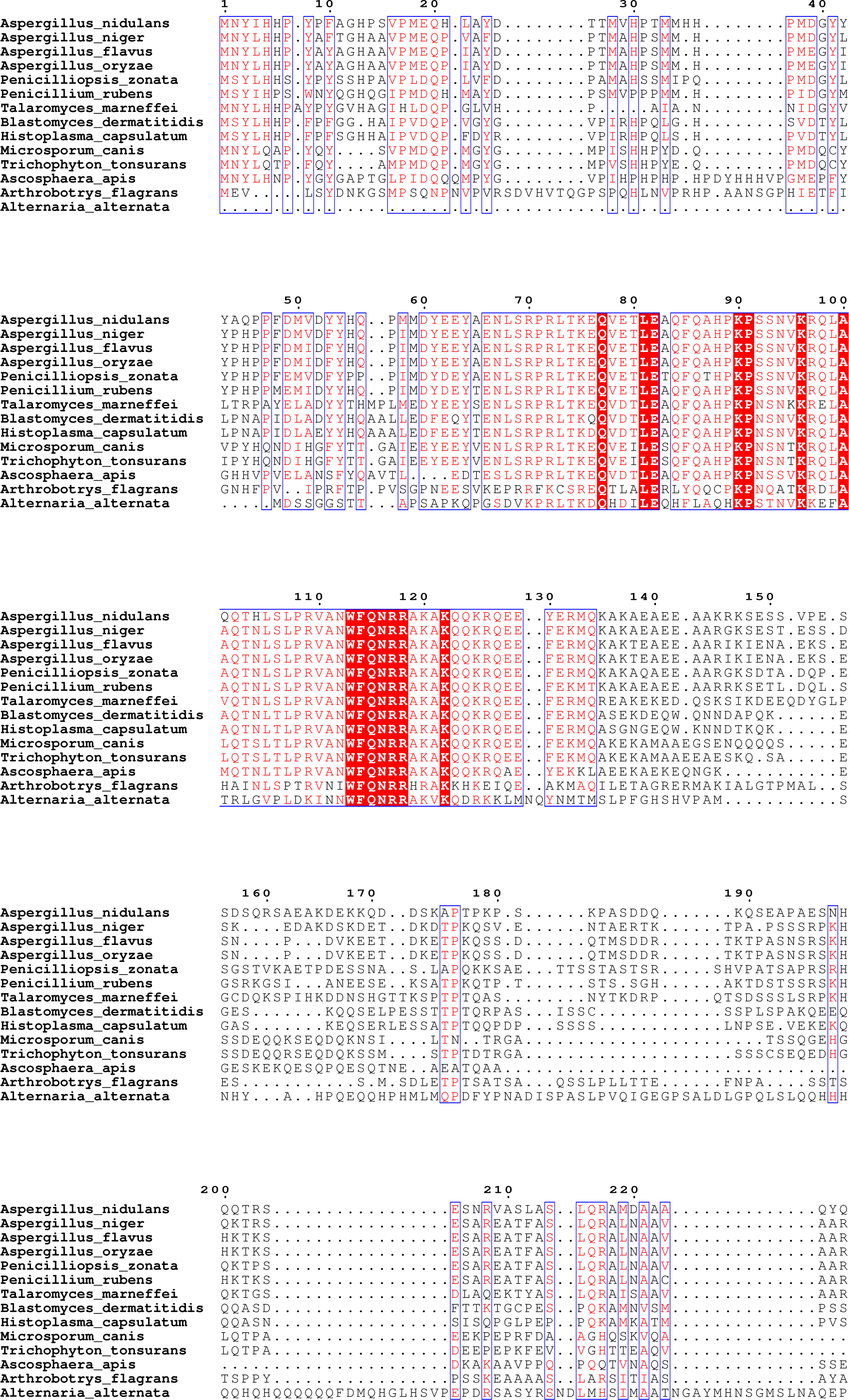
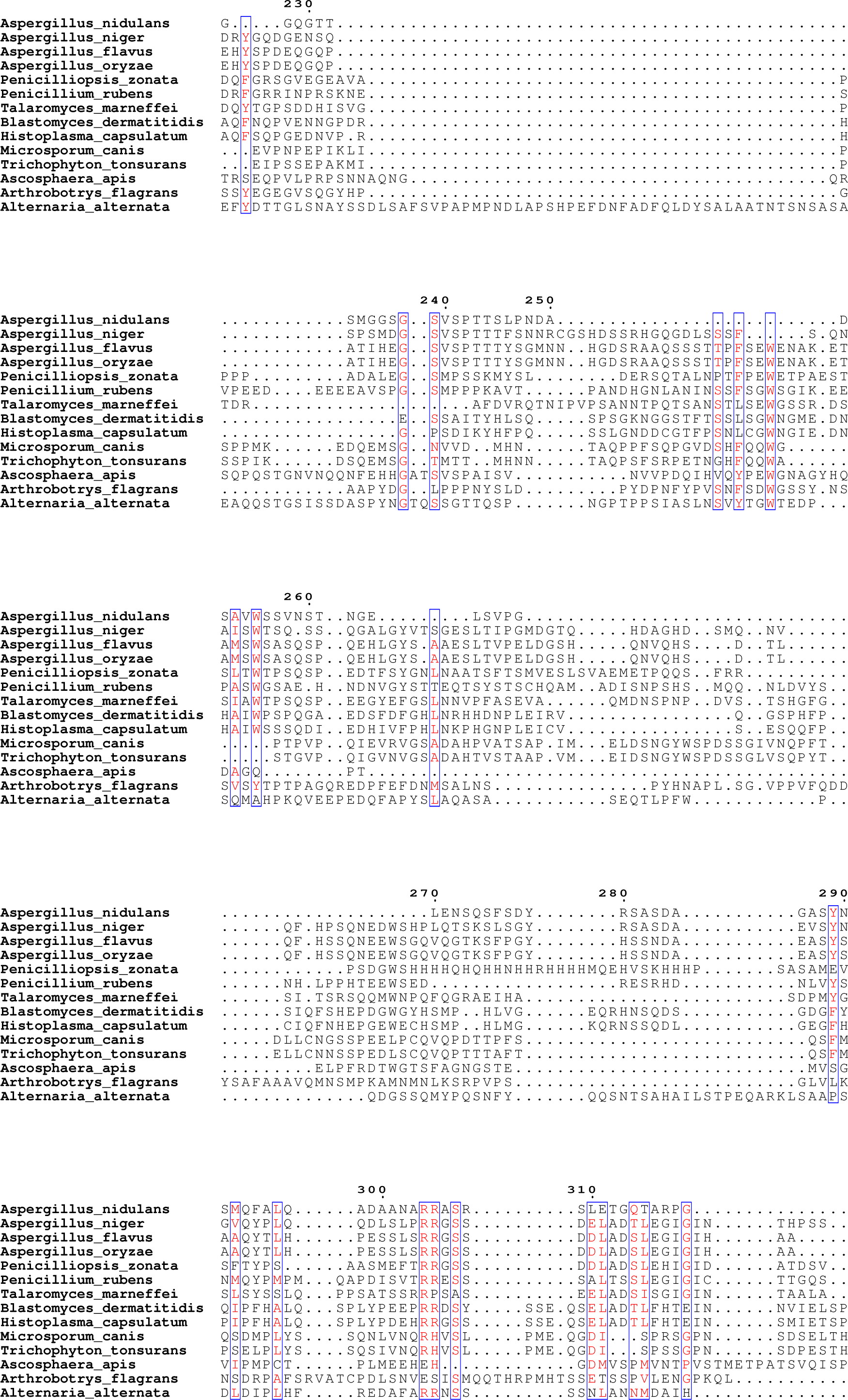
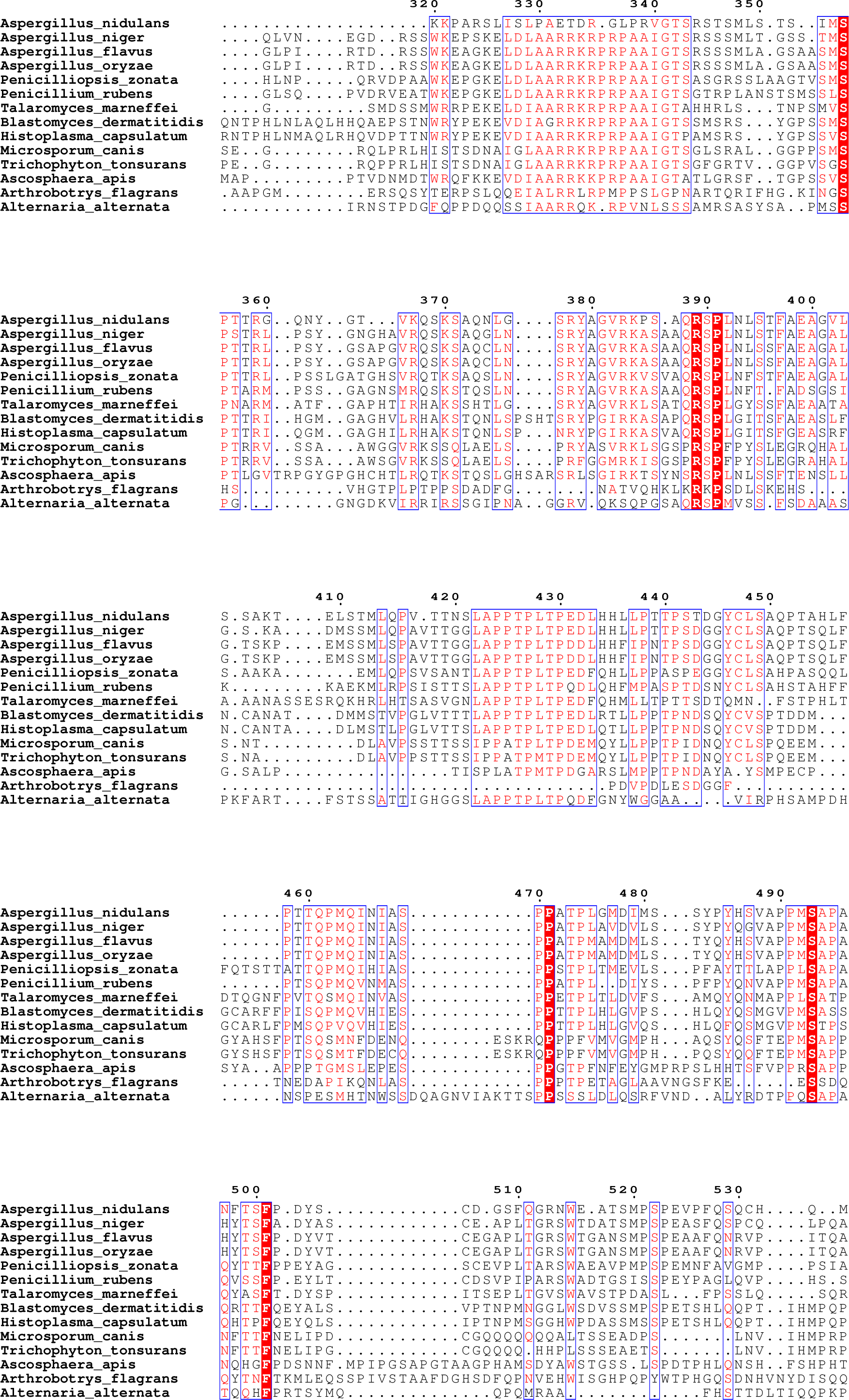
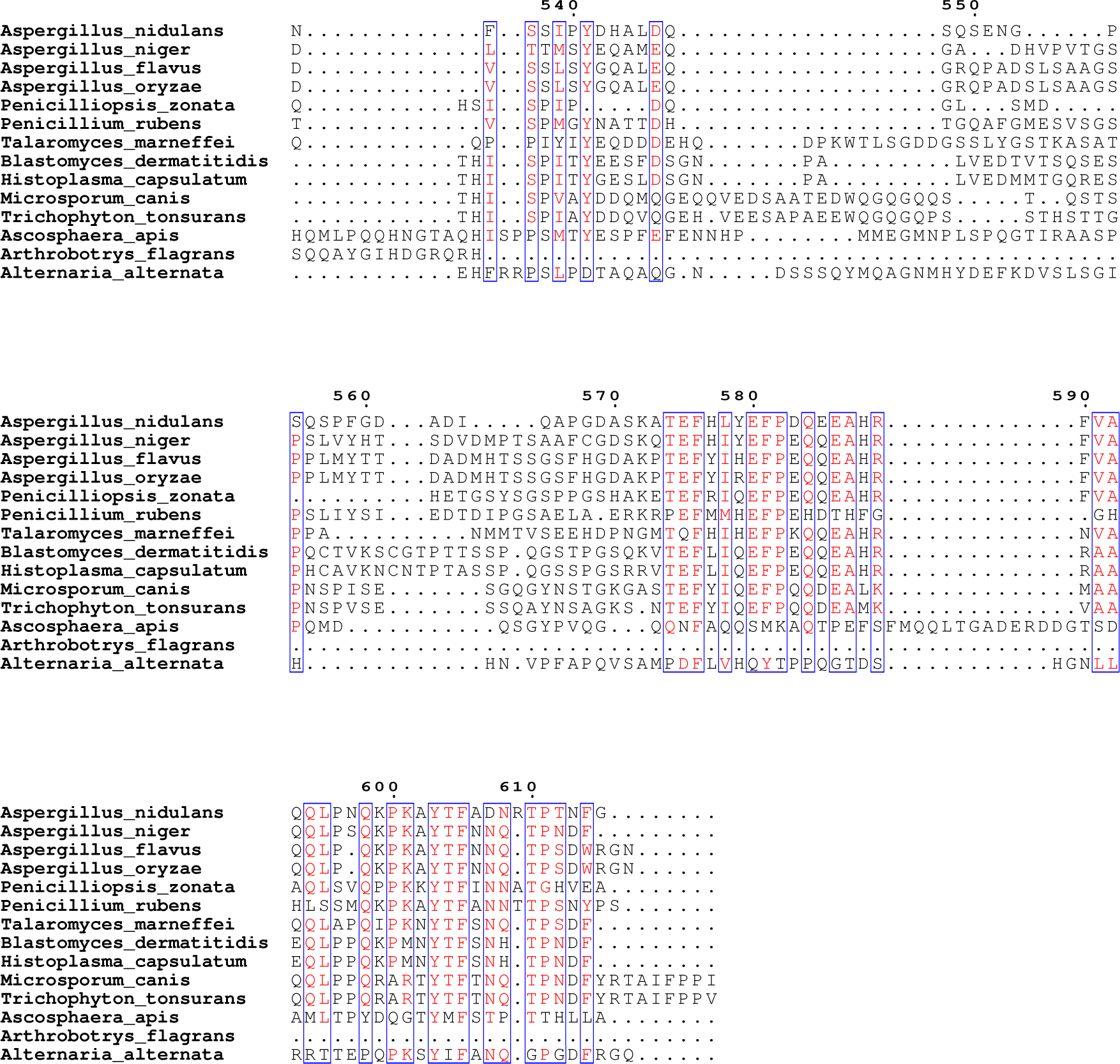
Multiple sequence alignment of *A. nidulans* HbxA with other fungal homologs. The HbxA deduced amino acid sequences were aligned using clustalOmega(https://www.ebi.ac.uk/Tools/msa/clustalo/). Data was visualized with boxshade using ENDscript server (https://espript.ibcp.fr/ESPript/cgi-bin/ESPript.cgi)(53)https://doi.org/10.1093/nar/gku316

**Table 3:**
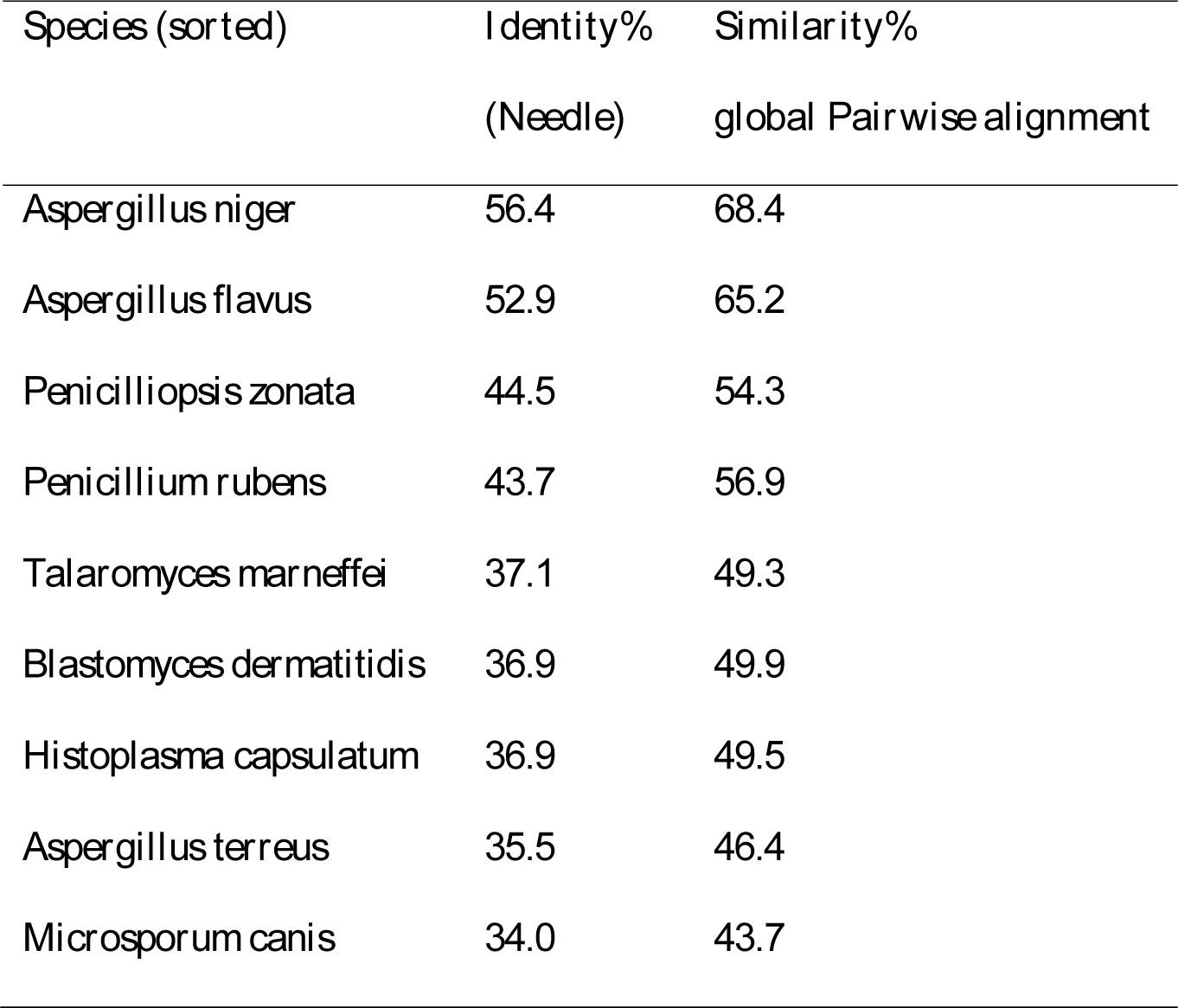
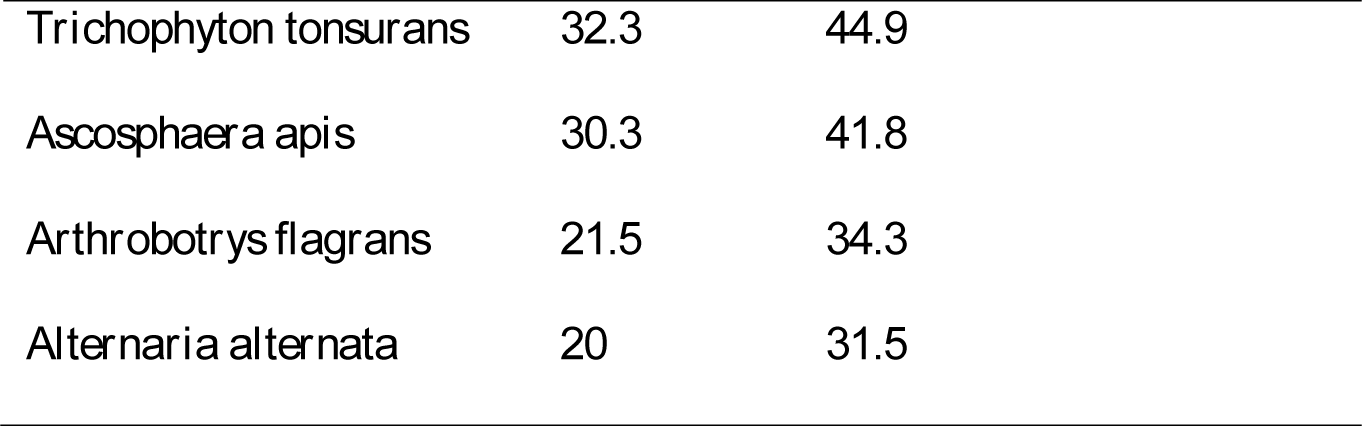
Phylogenetic analysis of *A. nidulans* HbxA and homologs in other fungal species. HbxA homologos were retrieved from FUNGIDB website and BLASTp was performed against protein sequence database. % similarity was found utilizing Pairwise sequence alignment using *A. nidulans* HbxA as search query against each protein of interest using EMBOSS Needle.

### hbxA is required for normal development in A. nidulans

To determine the regulatory scope of *hbxA* in *A. nidulans*, three strains were generated, a deletion strain, Δ*hbxA*, a complementation strain, *hbxA-*com, and an over-expression strain, OE*hbxA* (Fig 3). Deletion, complementation and overexpression strains were confirmed by diagnostic PCR, yielding the expected 3.01 kb PCR product for Δ*hbxA*, a 3.96 kb DNA fragment for *hbxA-*com and a 3.16 kb DNA fragment for OE*hbxA*. Our results confirmed that absence of *hbxA* results in a drastic reduction of conidiation (Fig 4), as previously shown(33,34).

**Fig 3:**
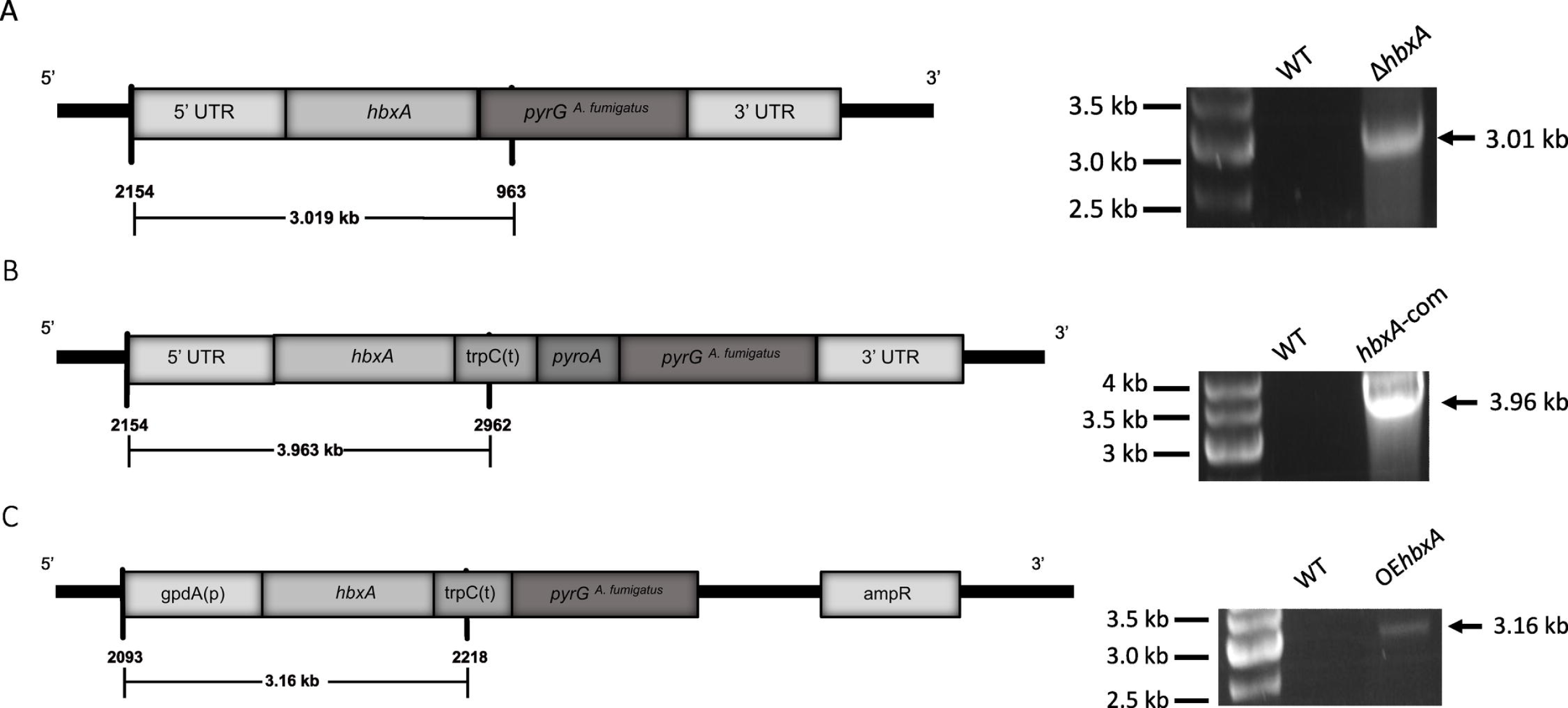
Generation of *A. nidulans hbxA* deletion, complementation and overexpression strains. Confirmation of the deletion (Δ*hbxA*), complementation (*hbxA*-com) and overexpression (OE*hbxA*) by diagnostic PCR. **(A)** The diagram shows replacement of *hbxA* with the marker gene *pyrG* by a double cross-over event. Primers P#2154/SD3 and P#963 were used for the diagnostic PCR, obtaining the predicted 3.01 kb product**. (B)** Schematic representation showing reintroduction of the wild-type *hbxA* allele at the *hbxA* locus in the deletion strain TSSP38.1. PCR with primers P#2154/SD3 and P#2962 confirmed the reintroduction of *hbxA* in the selected deletion strain; the expected 3.96 kb product was obtained. **(C)** Linear diagram of *hbxA* overexpression plasmid pSSP34.1. The overexpression transformant was confirmed by PCR with primers 2093 and 2218, which yielded the predicted 3.16 kb product.

**Fig 4.**
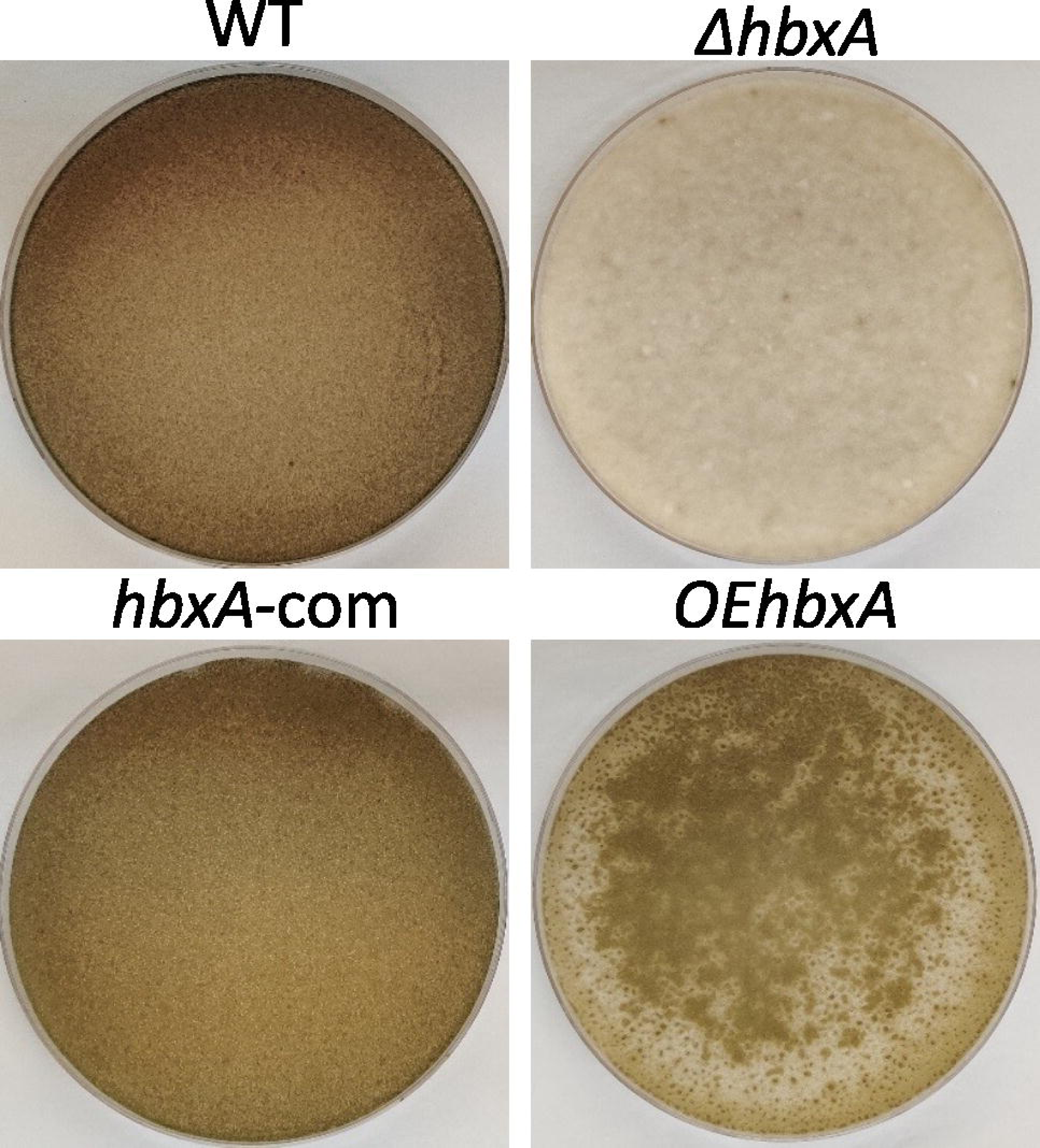
*hbxA* is required for normal conidiation in *A. nidulans*. Cultures of wild type, deletion, complementation and overexpression *hbxA* strains, top-agar inoculated on GMM and incubated for 7 days in the dark at 37°C.

### ShbxA regulates secondary metabolism

Our TLC analysis indicated that deletion of *hbxA* reduces sterigmatocystin (ST) production in *A. nidulans* by approximately 50 % when compared with levels in the wild-type strain (Fig 5).

**Fig 5.**
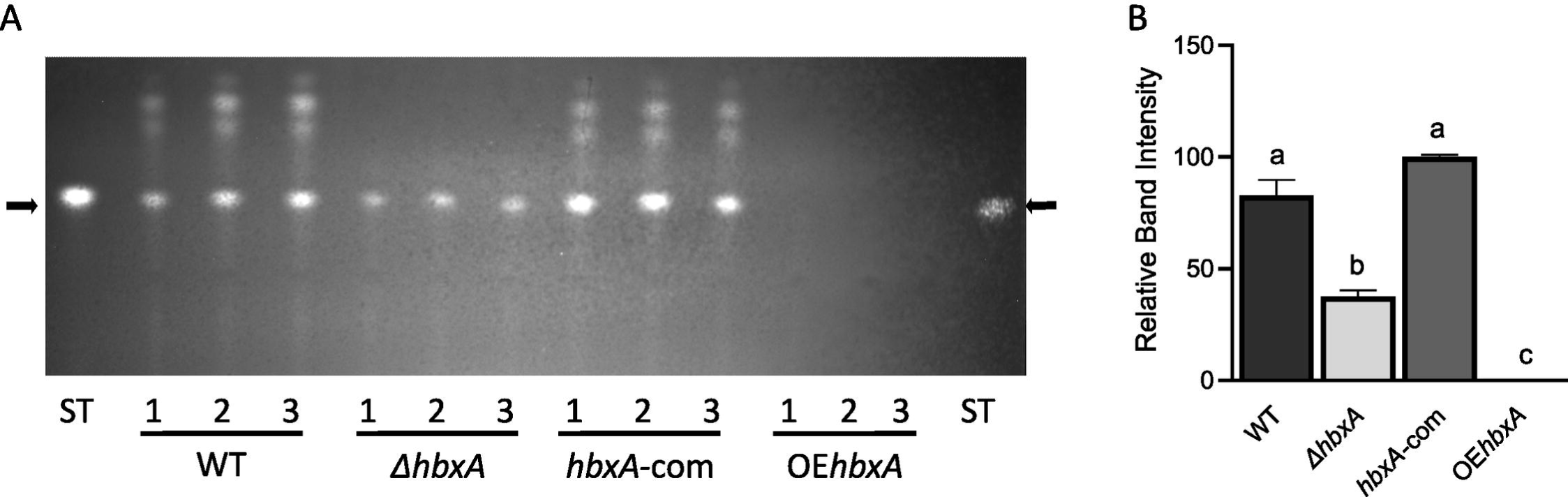
Effect of *hbxA* on the production of ST and other secondary metabolites in *A. nidulans.* Wild type, deletion, complementation and overexpression *hbxA* strains were top-agar inoculated on glucose minimum medium (GMM) and incubated for 3 days in the dark. **(A)** Extracts were analyzed by TLC. Black arrows indicate ST standard. The experiment was carried out with three replicates. **(B)** Densitometry of TLC analysis of ST levels. The desitometry was performed using the http://biochemlabsolutions.com/GelQuantNET.html website. Error bars represent the standard error. Columns of different letters represent values that are statistically different *p* value of <0.05

Importantly, overexpression of *hbxA* completely blocked ST production. Additionally, synthesis of other metabolites was also affected by deletion or forced overexpression of *hbxA* compared to the control strain. The absence of metabolites was particularly notable in the OE*hbxA* strain extracts. These results suggested that the regulatory role of *hbxA* is broader than originally expected, controlling not only developmental processes but also acting as a global regulator of secondary metabolism.

### hbxA*-*dependent transcriptome in A. nidulans

#### More than one thousand genes are regulated by hbxA in A. nidulans

RNA-sequencing analysis revealed that of the predicted 11286 genes present in *A. nidulans* genome (54), 552 were downregulated, and 195 were upregulated in the Δ*hbxA* strain compared with the wild-type control strain (Table 4, Fig 6). Over-expression of *hbxA* resulted in an even more pronounced effect on the *A. nidulans* transcriptome, where 1044 genes were downregulated, and 424 genes were upregulated in the OE*hbxA* strain in comparison to the wild type. In strong contrast, the comparison of the complementation strain and wild type showed that the two strains present very similar expression patterns. Expression of 618 genes in the *A. nidulans* genome was altered by either deletion or overexpression of *hbxA*, many of them presenting the same expression pattern of upregulation or downregulation when *hbxA* was either deleted or overexpressed (Fig 6).

**Fig 6.**
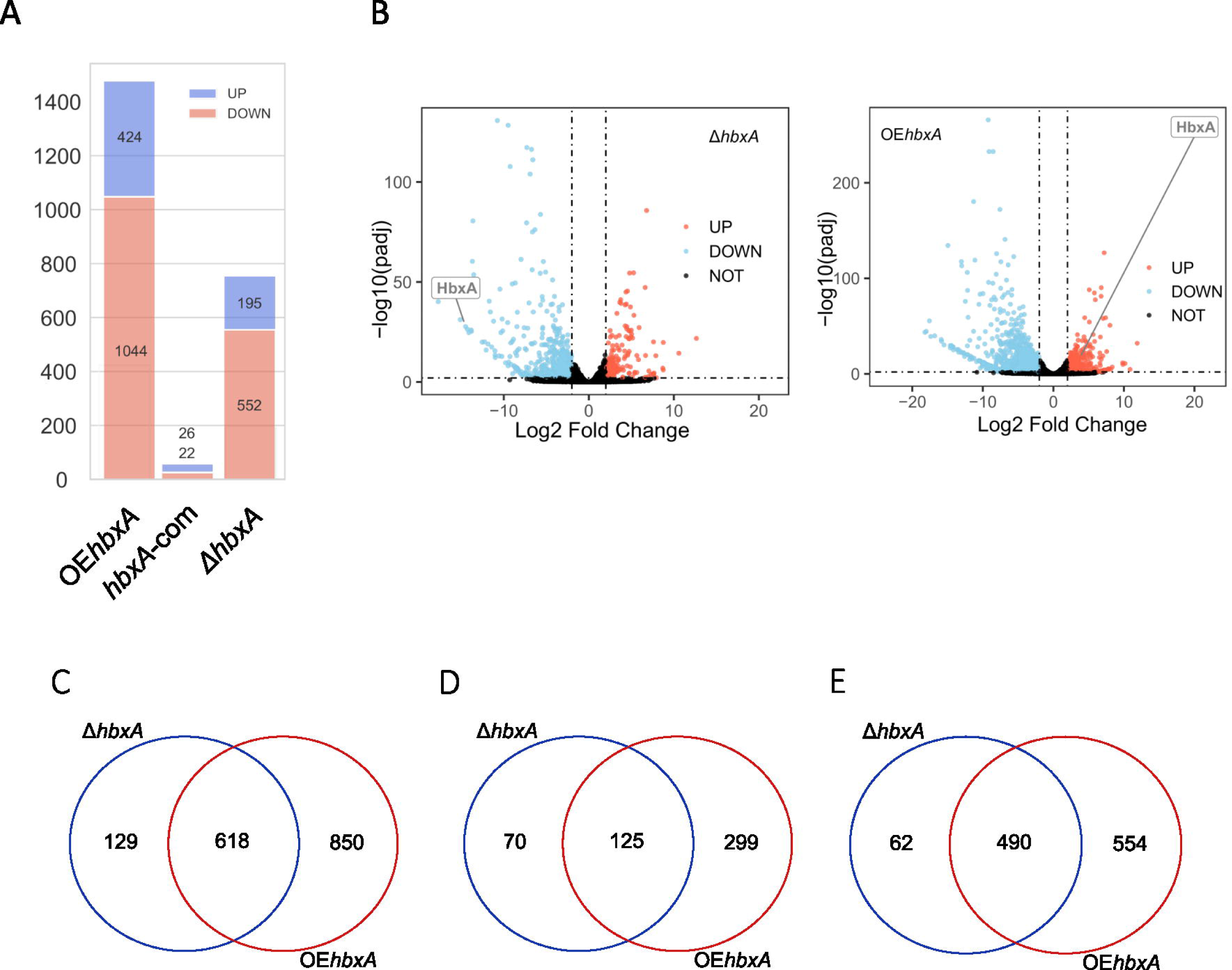
Number of DEGs in *A. nidulans* when expression of *hbxA* is altered by *hbxA* deletion or overexpression. (**A)** Number of significantly upregulated (purple) and significantly downregulated (orange) DEGs estimated by DeSeq2. **(B)** Volcano plot of log2 fold change vs. −log10 q-value of all the genes in Δ*hbxA*, and OE*hbxA* vs. control. Significantly upregulated genes are shown as red dots, significant down regulated genes are shown as blue dot and other genes are shown as black. The x-axis represents the log2 of the fold change determined by DeSeq2. The y-axis is the log10 of the adjusted p-value from DeSeq2. The cut offlog10 fold change value to determine the upregulated expression is greater than 2 while −2 is for down regulated expression. The −log10 q-value cutoff was set to 2 to determine the significant expression or not. **(C-D)** Venn Diagrams showing the overlap of DEGs in Δ*hbxA* and OE*hbxA* **(C)**, and the overlap of upregulated **(D)** and downregulated DEGs **(E)** in Δ*hbxA* and OE*hbxA*. Venn Diagrams were constructed using https://bioinformatics.psb.ugent.be/cgi-bin/liste/Venn/calculate_venn.htpl website.

#### Comparison of hbxA/hbx1 DEGs in A. nidulans and A. flavus

The comparison of the current *A. nidulans hbxA*-dependent transcriptome study with the previous *A. flavus hbx1* results (27) is shown in Fig 7. Only a small percentage of homologs were differentially expressed in the absence of *hbxA* and *hbx1* in *A. nidulans* and *A. flavus*, respectively, with respect to the corresponding wild types. Most of the DEGs in *A. nidulans* are not DEGs in the *A. flavus* study.

**Fig 7:**
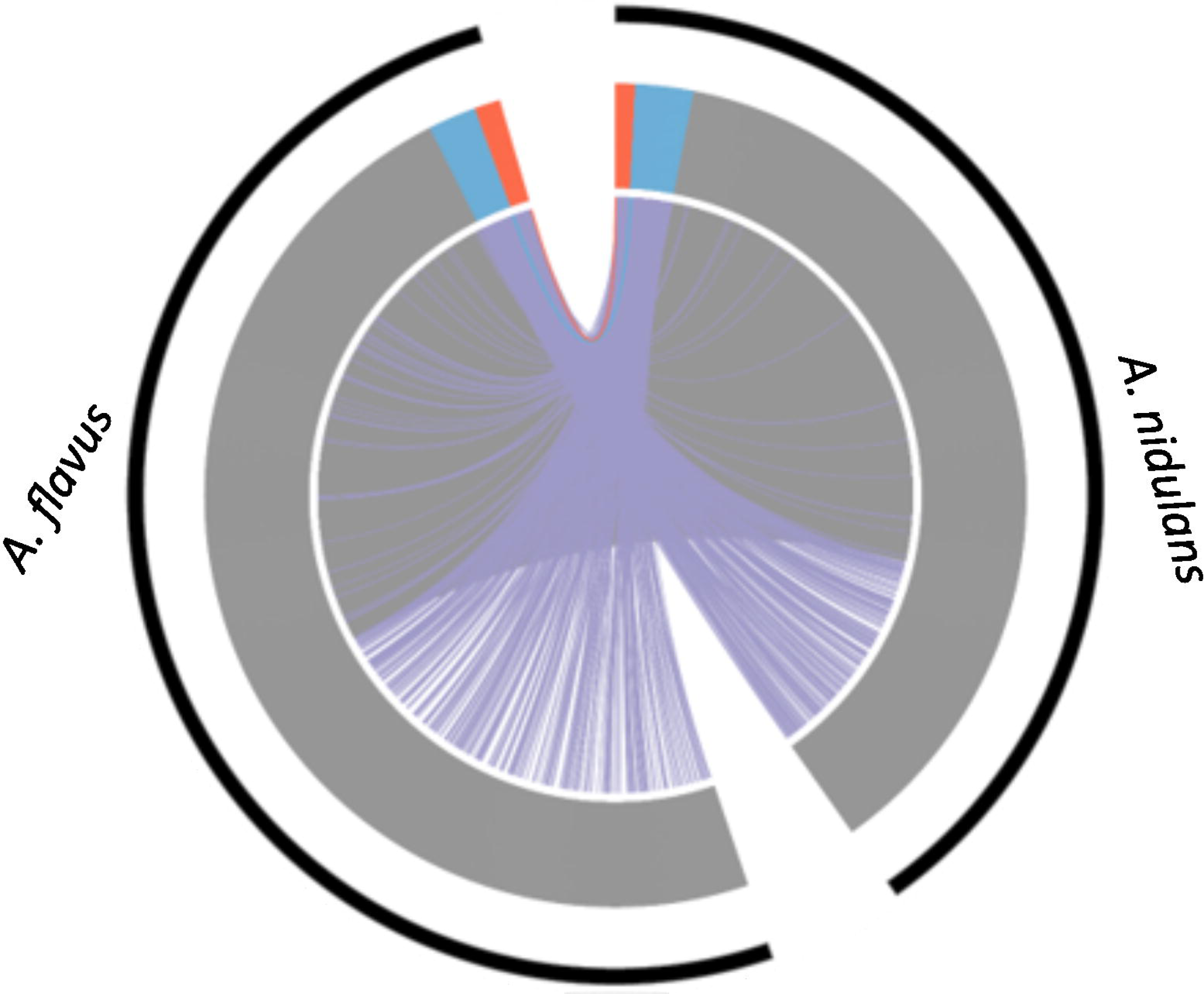
Comparison of orthologous genes affected by deletion of *hbxA* in *A. nidulans* and *A. flavus*. Both upregulated orthologous genes were colored in red. Both downregulated orthologous genes were colored in blue. No expression changed orthologous genes are colored in grey. Two orthologous genes having different regulation status are colored in purple. The significantly regulated genes were defined as |log2 fold change| <= 2 and q-value <= 0.05.

#### Expression of numerous TF genes is hbxA-dependent in A. nidulans

Based on our analysis, 74 out of 521 TFs genes in *A. nidulans* (51) were regulated by *hbxA* under the culture conditions assayed (Table 5). Some of these differentially expressed TF genes also presented the same expression pattern of upregulation or downregulation when *hbxA* was either deleted or overexpressed (Fig 8).

**Fig 8.**
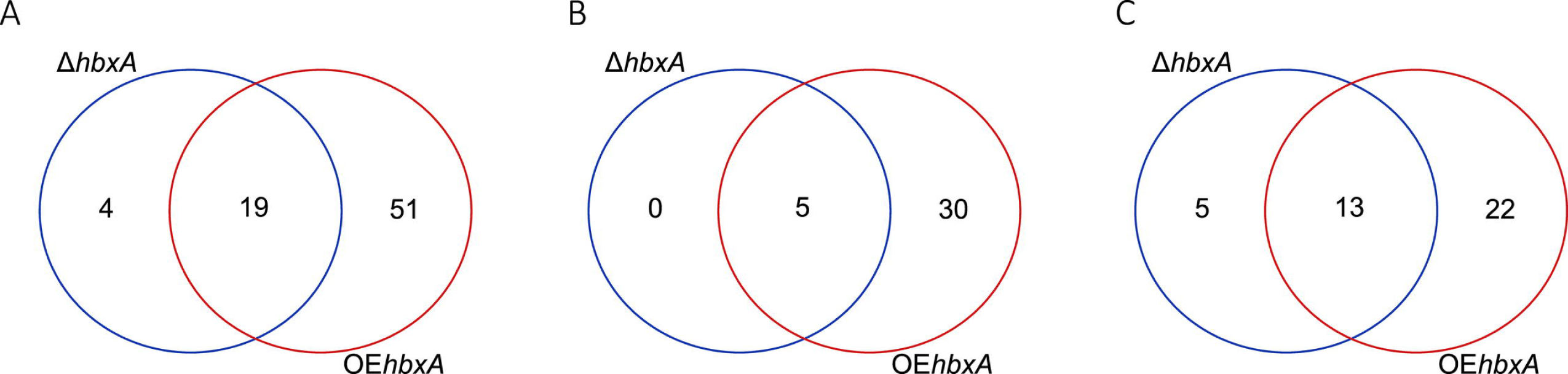
Number of transcription factor (TF) genes controlled by *hbxA in A. nidulans*. **(A)** Venn Diagram showing the overlap of differentially expressed TF genes in Δ*hbxA* and OE*hbxA.* **(B-C)** Venn Diagrams showing the overlap of upregulated **(B)** or downregulated **(C)** TF genes in Δ*hbxA* and OE*hbxA*. Venn Diagrams were constructed using https://bioinformatics.psb.ugent.be/cgi-bin/liste/Venn/calculate_venn.htpl website.

Our results indicated that overexpression of *hbxA* caused upregulation of developmental regulators, including genes of the central developmental pathway, *brlA*, *abaA* (55–58), *fluffy* genes *flbC* and *flbD* (59), and another HD-TF gene, *hbxB,* that regulates asexual and sexual development in *A. nidulans*(34). In addition, the developmental regulatory gene *zcfA* (60) was also upregulated by *hbxA* overexpression. Some of the upregulated TFs genes in OE*hbxA* are involved in both governing development as well as SM, such as the master transcription factor *mtfA* (37,61,62), *urdA,* (63), *sclB* (64), *osaA* (65), and *velB* (66). Other upregulated *hbxA*-dependent TF-DEGs annotated to be putatively involved in SM regulation include AN8391 and AN6788. Other upregulated TF genes have an important role in primary metabolism, such as *glcD*, which has a putative role in protein dimerization and activation of *areB*, (67), *galR,* which is known to regulate the D-galactose catabolic pathway (68) and *creA* repressor of carbon catabolite (69). Other upregulated TF genes were *rfeC,* whose ortholog in *Saccharomyces cerevisiae* promotes *FLO11* expression (70), the *mcnB* fork-head like transcription factor (71), as well as expression of some other uncharacterized putative transcription factors genes (Table 5).

Overexpression of *hbxA* in *A. nidulans* caused downregulation of other developmental genes such as *fhpA*, with a role in sexual development (72), *mat1*, involved in activation of the alpha-domain mating-type protein (73). Overexpression of *hbxA* also caused downregulation of *metZ*, a transcription factor involved in the regulation of sulfur metabolism (74). TF genes AN8377, AN8645, AN3385 and AN8918 predicted to be involved in SM, are also downregulated in this strain (Table S2).

Interestingly, deletion of *hbxA* also resulted in an increase in the expression of *brlA*, *abaA,* and *urdA*, as in the case of OE*hbxA.* It also increased the expression of *tah-3,* which is involved in conidiophore development and tolerance for harsh plasma environment (75) (Table 5). Deletion of *hbxA* also upregulated *veA* (Table 4). The *veA* gene product, VeA, which contains a NF-κ-B like DNA-binding domain (76), is well known as a global regular that interacts with at least nine other proteins, LlmF, VapA, VipA, VipC, VelB, MpkB, FphA, LreB and LaeA (77), governing several signaling pathways and consequently multiple cellular processes, including development and SM (25).

Absence of *hbxA* in *A. nidulans* downregulated the expression of various transcription factors, including the gene encoding the alpha-domain mating-type protein, *mat1*(73), as in overexpression of *hbxA*. Deletion of *hbxA* also showed downregulation of *metZ,* involved in methionine biosynthesis (78) the nitrogen-dependent *mdpE*, which regulates production of a secondary metabolite called monodictyphenone (79). The putative SM TF gene AN4933 is downregulated, and AN3385, AN8645 and AN8918 are also downregulated in deletion *hbxA*, as in OE*hbxA*.

#### hbxA affects the expression of genes in SM gene clusters and biosynthesis of natural products in A. nidulans

Our TLC analysis revealed that both deletion and overexpression of *hbxA* negatively affect ST production (Fig 5) as well as the production of other secondary metabolites. Furthermore, FunCat enrichment analysis revealed that differentially regulated genes in the Δ*hbxA* versus wild type and OEveA versus wild type comparisons have significant functional overlap (Fig 9). DEGs genes are dramatically enriched for secondary metabolism-related processes for both; most of those genes are downregulated when *hbxA* is either deleted or overexpressed, particularly in the latter. Other categories showing enrichment include disease, virulence, and defense; virulence disease factors; C-compound and carbohydrate metabolism; and detoxification.

**Fig 9.**
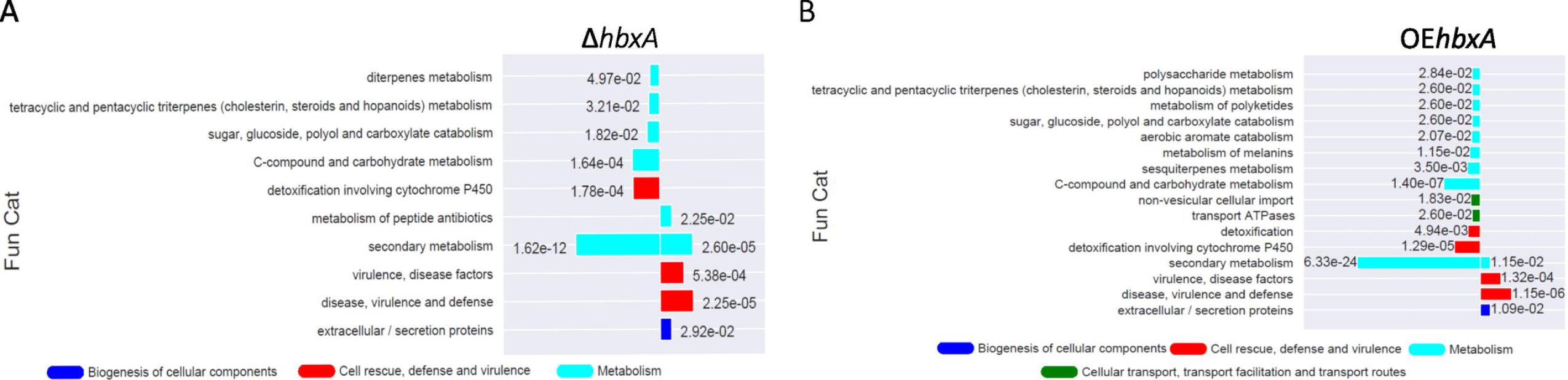
FunCat enrichment of significant DEGs found in (A) Δ***hbxA* and (B) OE*hbxA* vs. control**. The −log10 of the q-value of DEGs in each term is proportional to the length of the bars. FunCat annotations and q-value is determined by FungiFun2 webserver. Downregulated genes are to the left of the origin and up regulated genes to the right.

To gain further understanding of the effect of *hbxA* on SM in *A. nidulans*, as part of our transcriptome analysis, we identified DEGs in SM gene clusters and analyzed concomitant production of secondary metabolites by a metabolomics approach. Our study revealed that production of nidulaninA, nidulanin B and nidulanin D are *hbxA*-dependent (Fig 10A-C). Both, deletion and overexpression of *hbxA,* completely inhibited the production of these compounds. In addition, the Heatmap shown in Fig 10D indicates downregulation of some of the genes in the nidulanin cluster (80), including the NRPS coding gene, *nlsA*, in both Δ*hbxA* and OE*hbxA*. This reduction in *nlsA* expression was particularly notable in the latter.

**Fig 10:**
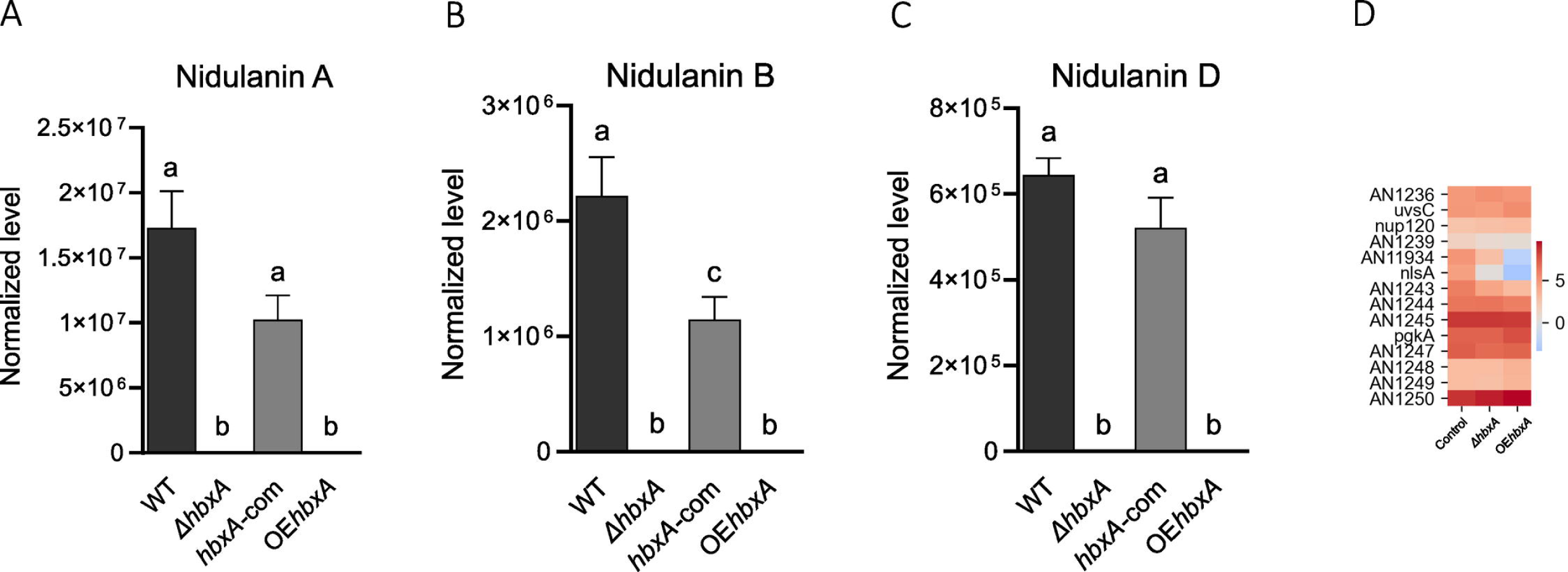
*hbxA* regulates the production of nidulanins in *A. nidulans.* Wild-type (WT), deletion (Δ*hbxA*), complementation (*hbxA-com*) and overexpression (OE*hbxA*) strains were top-agar inoculated on solid glucose minimum medium (GMM) at 37°C for 72 h, when samples were collected, extracted with chloroform and analyzed by LC-HRMS in positive mode **(A-C)** Quantification of nidulanin A (*m*/*z* 604.34943), B (*m*/*z* 620.34404) , D (*m*/*z* 536.28659) respectively. **(D)** Heat map of TPM values of nidulanin cluster (DEGs) expression in *A. nidulans* Δ*hbxA* and OE*hbxA* with respect to wild type strain on a log scale found in Inglis et al.(50). The TPM value of each gene was calculated by averaging all the TPM values of all replicates.

LC-MS analysis of ST confirmed the TLC results, indicating that production of this mycotoxin was reduced in Δ*hbxA* and absent in the overexpression strains (Fig 11). Unexpectedly, the Heatmap in Fig 11B shows that most of the ST genes were not downregulated in the deletion strain with respect to the wild type. However, most of the genes in this cluster were downregulated in the overexpression strain, excluding the structural genes *stcK, stcJ, stcF* and *stcC,* and the regulator, *aflR*.

**Fig 11.**
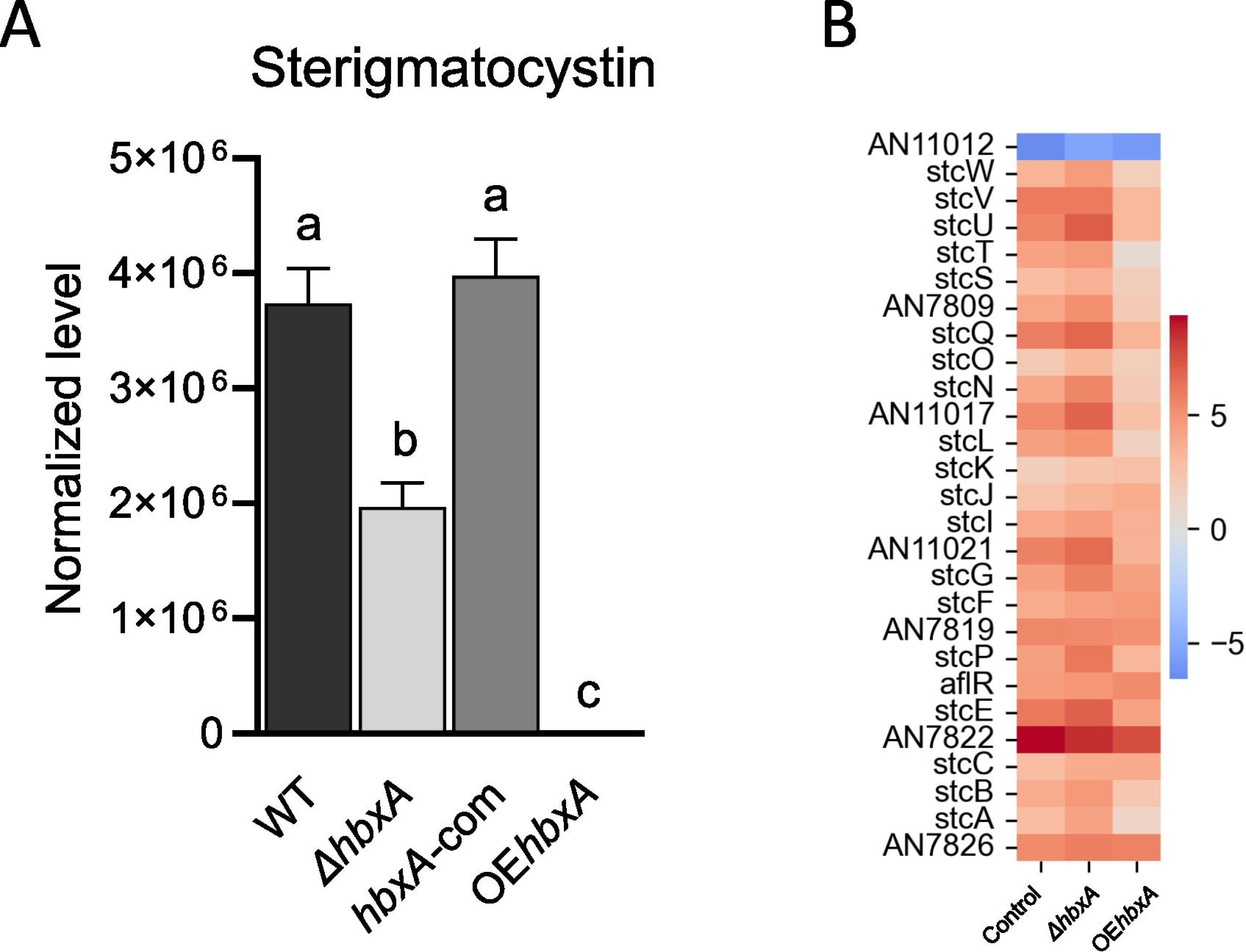
*hbxA* regulates the production of ST in *A. nidulans.* Wild type (WT), deletion (Δ*hbxA*), complementation (*hbxA-com*) and overexpression (OE*hbxA*) strains were top-agar inoculated on solid glucose minimum medium (GMM) at 37°C for 72 h, when samples were collected, extracted with chloroform and analyzed by LC-HRMS in positive mode. **(A)** Quantification of ST (*m*/*z* 325.07014). **(B)** Heat map of TPM values of ST cluster (DEGs) expression in *A. nidulans* Δ*hbxA* and OE*hbxA* with respect to wild type strain on a log scale found in Inglis et al. (50). The TPM value of each gene was calculated by averaging all the TPM values of all replicates.

In addition, both Δ*hbxA* and OE*hbxA* strains were unable to synthesize the meroterpenoids austinol and dehydroaustinol under conditions conducive to their production in the wild type (Fig 12). The genes involved in the synthesis of these two compounds are grouped in two clusters, A and B (81). Our transcriptome analysis revealed that most of the genes in these two clusters are downregulated in the *hbxA* deletion and also in the overexpression strains compared to the control (Fig 12C and D). For example, genes *ausA-D* are down regulated in both Δ*hbxA* and OE*hbxA* in gene cluster A. In cluster B, genes *ausE-G* and *ausM* are also downregulated in Δ*hbxA* and OE*hbxA*. Additionally, expression of *ausH*. *ausL* and *ausN* is reduced in OE*hbxA* with respect to the wild type.

**Fig 12.**
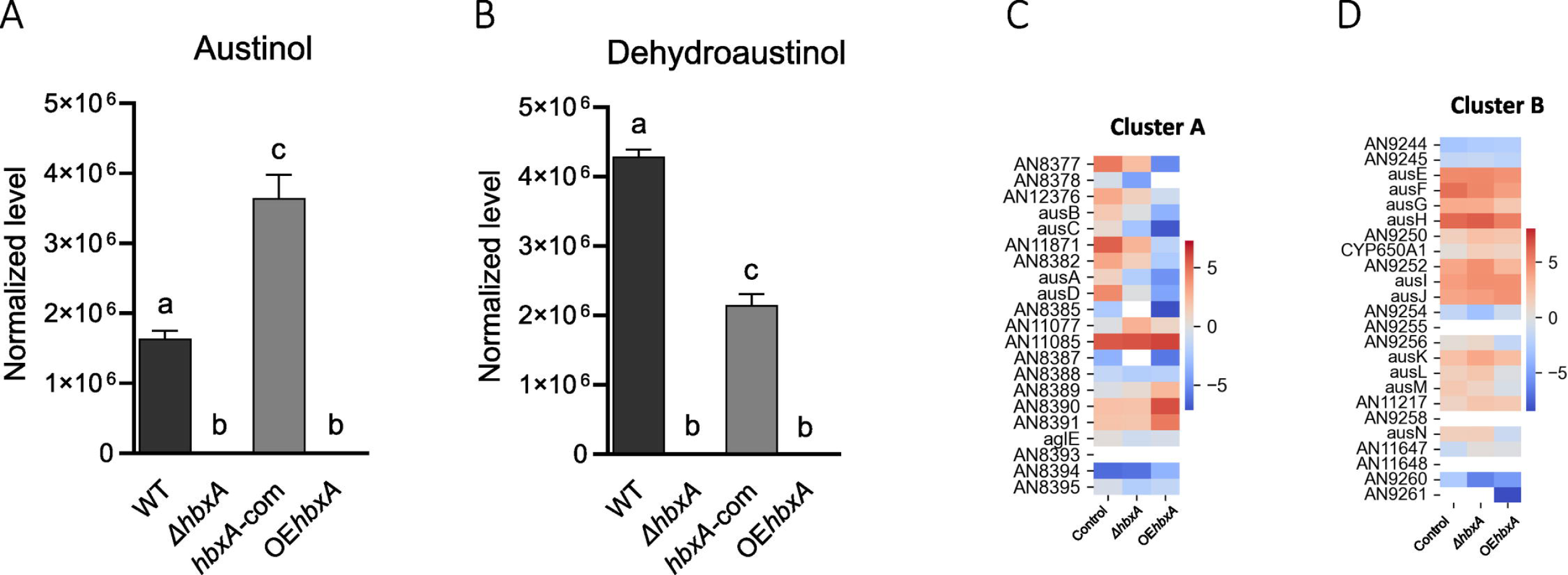
*hbxA* regulates the production of austinol and dehydroaustinol in *A. nidulans.* Wild type (WT), deletion (Δ*hbxA*), complementation (*hbxA-com*), and overexpression (OE*hbxA*) strains were top-agar inoculated on solid glucose minimum medium (GMM) at 37°C for 72 h, when samples were collected, extracted with chloroform and analyzed by LC-HRMS in positive mode. Quantification of **(A)** austinol (*m*/*z* 459.20059) and **(B)** dehydroaustinol (*m*/*z* 457.18524) compounds by full MS spectra resolution of 60,000 with a range of mass-to-charge ratio (*m/z*) set to 50LtoL800. **(C & D)** Heatmap of TPM values of austinol cluster (DEGs) expression in *A. nidulans* Δ*hbxA and* OE*hbxA* with respect to wild type strain on a log scale found in Inglis et al. (50). The TPM value of each gene was calculated by averaging all the TPM values of all replicates.

Our metabolomics study also indicated that the production of three novel, unknown secondary metabolites was altered when *hbxA* was not expressed at wild-type levels. Two of these compounds (m/z 423 and m/z 518 observed in negative mode) were absent in the *hbxA* deletion strain and also in the overexpression strain (Fig 13). The third novel compound (m/z 489 in negative mode) was produced at remarkably high levels in the *hbxA* deletion strain compared to those in the wild type (Fig 13B).

**Fig 13:**
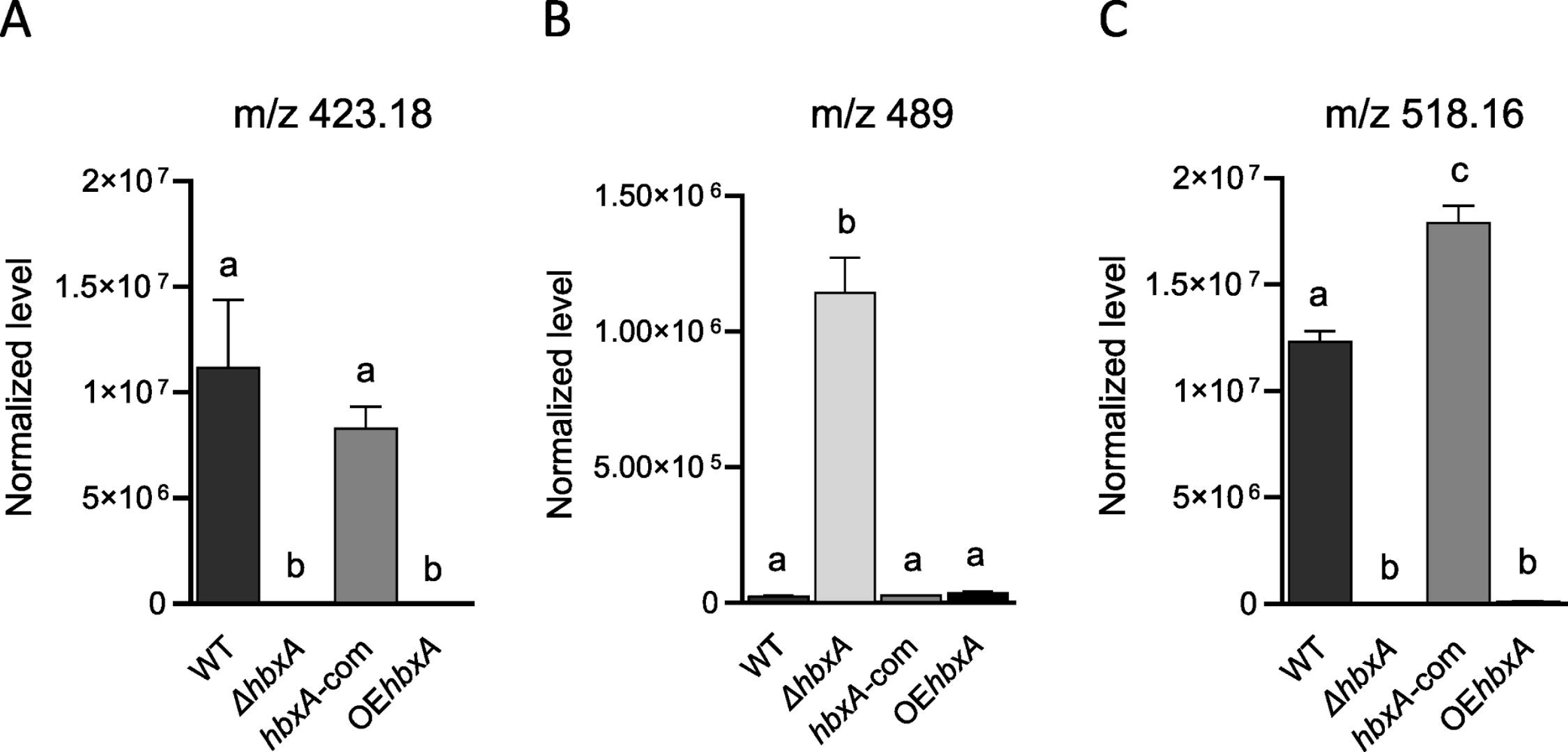
*hbxA* regulates the production of novel uncharacterized metabolites in *A. nidulans.* Wild type (WT), deletion (Δ*hbxA*), complementation (*hbxA-com*), and overexpression (OE*hbxA*) strains were top-agar inoculated on solid glucose minimum medium (GMM) at 37 °C for 72 h, when samples were collected, extracted with chloroform and analyzed by LC-HRMS in negative mode. **(A-C)** Quantification of novel uncharacterized metabolites with *m*/*z* of 423.18012, 489.18082, and 518.16482, respectively.

## DISCUSSION

HD-TFs have been shown to govern development in eukaryotes (13–15), including fungi (13,14,16–21). Previous reports, together with the present study, indicate that these regulators are conserved across different fungal genera. (Zheng et al., 2012; Ghosh et al., 2015). In *A. flavus*, *hbx1*, an ortholog of *hbxA*, is also required for developmental processes, regulating genes in the conidiation central pathway, such as *brlA* and *wetA* (17,27) as well as *flbA, flbC, flbD, flbE, fluG* and *mat1-1*(27). In *A. fumigatus*, *hbxA* promotes *brlA*, *abaA* and *wetA*, as well as *flbB*, *flbD* and *fluG* expression(28). Similarly, *hbxA* regulates conidiation in *A. nidulans*(33,34); our transcriptome study showed that *hbxA* not only regulates *brlA,* as shown in (34), but also *abaA*, *flbC* and *flbD*. These studies support that the *hbxA*-dependent regulatory mechanism of conidiation is at least in part conserved in these three Aspergillus species and possibly in other species of this genus.

Interestingly, our results revealed a broader regulatory scope for *hbxA* in *A. nidulans*, with more than one thousand DEGs when *hbxA* was deleted or overexpressed in this model organism, including numerous transcription factor genes. This was also the case for *A. flavus hbx1*(27). However, most of the DEGs in *A. nidulans* are not DEGs in *A. flavus*; only a small percentage of homologs where DEGs in the *hbxA* and *hbx1* mutants with respect to the controls. This suggests that although the conservation of some of the regulatory mechanisms controlling conidiation appears conserved, a great part of its regulatory input is specialized in different fungal species.

Some of the TF genes involved in governing development that were found *hbxA*-dependent also control secondary metabolism in *A. nidulans*, for example, *mtfA* (37,61,62), *urdA* (63)*, sclB* (64), *osaA* (65) and *velB* (66). Furthermore, FunCat functional enrichment analysis showed that the category of secondary metabolism-related processes was, by far, the most enriched in *A. nidulans*. Our study showed that in *A. nidulans,* numerous genes in SM gene clusters were regulated by *hbxA.* The secondary metabolism category was also enriched in *A. flavus* (27)However, the wide variation of biosynthetic gene clusters across fungal species, even in those phylogenetically close (82) could explain that although the major functional category is the same in both species, namely SM, the percentage of differentially expressed homologs is low. For example, *A. flavus hbx1* regulates genes in the aflatoxin, cyclopiazonic acid, aflatrem, asparasone, piperazine, and aflavarin gene clusters(27), while in *A. nidulans,* our study shows that *hbxA* controls genes in other gene clusters such as those responsible for the synthesis of nidulanins A, B and D, austinol and dehydroaustinol. *A. nidulans* HbxA also control genes in the ST gene cluster, which is partially conserved with that of aflatoxin in *A. flavus*. The regulatory pattern was similar; absence of both *hbxA* and *hbx1* resulted in a reduction of toxin production (17,27). In *A. flavus* deletion of *hbx1* downregulated *aflR* and other genes in the aflatoxin gene cluster. However, this was not the case in *A. nidulans*, suggesting that the lower levels of ST in the deletion strain, verified by both TLC and LC-MS, could be due to other factor(s). Our study showed *veA* expression is *hbxA*-dependent. VeA is a global regulator that orchestrates numerous biological processes in fungi (25,26), such as development and SM. VeA has been shown to regulate the production of aflatoxisomes in *A. parasiticus* (83). It is possible that *hbxA*, in a *veA*-dependent manner, could also influence compartmentalization of ST production in *A. nidulans*. This reduction in ST in the deletion strain, contrast with a previous report (34) where an increase in ST was described. It is possible that different experimental conditions in both studies could have resulted in different outcomes. Nevertheless, the most striking result is the effect of *hbxA* overexpression on ST biosynthesis as well as on the production of other metabolites. The complete elimination of ST production by *hbxA* overexpression was, in this case, accompanied by the downregulation of genes in the ST gene cluster. However, this downregulation of ST genes was, as in the case of the deletion strain, not mediated by changes in *aflR* expression.

Our study revealed that *hbxA* regulates key genes in the nidulanin gene cluster and, consequently, affects the production of the cyclic tetrapeptides nidulinins A, B and D. These compound are found in *Aspergillus* and *Penicillium* species. The function of nidulanins is not yet known. As in the case of ST, both deletion or overexpression of *hbxA* resulted in reduction or elimination of nidulinins A, B and D production, suggesting that, as in the case of VeA, certain balanced stoichiometry with respect to other regulatory factors could be needed for proper function, perhaps also interacting with other regulatory proteins. One of the genes downregulated in both deletion *hbxA* and overexpression *hbxA* strains is *nlsA*, encoding a non-ribosomal peptide synthase necessary for the synthesis of nidulanin. This enzyme has been shown to also be involved in the synthesis of fungisporin (84), which presents antibacterial activity (85), however fungisporin was not detected in our study under the conditions tested.

LC-MS indicated that *hbxA* also controls austinol and dehydroaustinol production. These are two meroterpenoids produced from polyketide and terpenoid precursors. Both austinol and dehydroaustinol have been shown to inhibit the neuraminidase enzyme, suggesting a potential for the development of new antiviral drugs (86). Austinol also showed antibacterial activity (87). Alteration of wild-type *hbxA* transcription by deletion or forced overexpression also resulted in a lack of production of these compounds, further supporting the possibility of a necessary stoichiometry with other regulatory partners. Two separate gene clusters, A and B (81,88), are required for the synthesis of these compounds. Both deletion and overexpression of *hbxA* showed profound changes in the expression profile of both gene clusters, with numerous downregulated structural genes, including the polyketide synthase gene *ausA.* The prenyltransferase gene *ausN* was also downregulated in the overexpression strain.

In addition, our metabolomics analysis indicated that *A. nidulans hbxA* also controls the production of three unknown novel compounds. Synthesis of two of these metabolites (m/z &423 and m/z 528) did not occur in the absence of *hbxA* or when this gene was overexpressed, while the third novel compound (m/z 489) was produced at strikingly high levels in the *hbxA* deletion strain. The identity, the association with MS gene clusters, or bioactive properties of these compounds are still known and will be the subject of future studies.

Regarding additional roles of *hbxA* in *A. nidulans*, besides those in development and SM, our FunCat functional enrichment analysis also indicated a possible role in primary metabolism, with enrichment in the carbon-compound and carbohydrate metabolism category, particularly in the *hbxA* overexpression strain. Upregulation of the carbon catabolite repressor TF gene *creA* (69) was observed in this strain. *creA* is also under *hbx1* regulation in *A. flavus*(27). Other *A. nidulans hbxA*-dependent regulatory genes involved in primary metabolism were, for example, *galR,* which regulates the D-galactose catabolic pathway (68), and *glcD*, which has a putative role in protein dimerization with and activation of *areB,* involved in nitrogen metabolism (67). Other enriched categories were detoxification, virulence and disease factors and defense, suggesting its possible involvement in pathogenesis. This agrees with the fact that the *hbxA* homolog in *A. fumigatus* was shown to affect virulence in *A. fumigatus* (28)

In conclusion, we have shown that the regulatory TF gene *hbxA* governs the expression of hundreds of genes in *A. nidulans*, modulating not only developmental genes, but also multiple regulatory pathways. Consequently, *hbxA* governs different important aspects of this fungus’ biology, including a remarkable role in SM, regulating expression of several SM gene clusters and natural product biosynthesis, including some novel compounds. Additionally, genes associated with other cellular processes such as primary metabolisms, as well as defense and virulence, are also influenced by *hbxA.* Interestingly, a functional conservation exists between *hbxA* homologs in other *Aspergillus* species and possibly in other fungi.

## ACKNOWLEDGEMENTS

This study was funded by Northern Illinois University

## Notes

### Competing Interest Statement

The authors have declared no competing interest.

## REFERENCES

1. Caesar LK, Kelleher NL, Keller NP. In the fungus where it happens: History and future propelling Aspergillus nidulans as the archetype of natural products research. Fungal Genet Biol. 2020 Nov 1;144:103477.

2. Oiartzabal-Arano E, Perez-de-Nanclares-Arregi E, Espeso EA, Etxebeste O. Apical control of conidiation in Aspergillus nidulans. Curr Genet. 2016 May 1;62(2):371–7.

3. Yu JH. Regulation of Development in Aspergillus nidulans and Aspergillus fumigatus. Mycobiology. 2010 Dec 31;38(4):229–37.

4. Etxebeste O, Garzia A, Espeso EA, Ugalde U. Aspergillus nidulans asexual development: making the most of cellular modules. Trends Microbiol. 2010 Dec;18(12):569–76.

5. Aramayo R, Timberlake W e. The Aspergillus nidulans yA gene is regulated by abaA. EMBO J. 1993 May;12(5):2039–48.

6. Hermann TE, Kurtz MB, Champe SP. Laccase localized in hulle cells and cleistothecial primordia of Aspergillus nidulans. J Bacteriol. 1983 May;154(2):955–64.

7. Bussink HJ, Osmani SA. A cyclin-dependent kinase family member (PHOA) is required to link developmental fate to environmental conditions in Aspergillus nidulans | The EMBO Journal . 1998. Available from: https://www.embopress.org/doi/full/10.1093/emboj/17.14.3990

8. Clutterbuck AJ. A mutational analysis of conidial development in Aspergillus nidulans. Genetics. 1969 Oct 1;63(2):317–27.

9. Han KH, Han KY, Yu JH, Chae KS, Jahng KY, Han DM. The nsdD gene encodes a putative GATA-type transcription factor necessary for sexual development of Aspergillus nidulans. Mol Microbiol. 2001 Jul 1;41(2):299–309.

10. Kirk KE, Morris NR. The tubB alpha-tubulin gene is essential for sexual development in Aspergillus nidulans. Genes Dev. 1991 Nov 1;5(11):2014–23.

11. Lee, Kim, S., Kim, S. J., Han, D. M., Jahng, K. Y., Chae, K. S. The lsdA gene is necessary for sexual development inhibition by a salt in Aspergillus nidulans | SpringerLink. 2001. Available from: https://link.springer.com/article/10.1007/s002940100206

12. Miller KY, Toennis TM, Adams TH, Miller BL. Isolation and transcriptional characterization of a morphological modifier: the Aspergillus nidulans stunted (stuA) gene. Mol Gen Genet MGG. 1991 Jun 1;227(2):285–92.

13. Holland PWH. Evolution of homeobox genes. Wiley Interdiscip Rev Dev Biol. 2013 Feb;2(1):31–45.

14. Mukherjee K, Brocchieri L, Bürglin TR. A comprehensive classification and evolutionary analysis of plant homeobox genes. Mol Biol Evol. 2009 Dec;26(12):2775–94.

15. Svingen T, Tonissen KF. Hox transcription factors and their elusive mammalian gene targets. Heredity. 2006 Aug;97(2):88–96.

16. Antal Z, Rascle C, Cimerman A, Viaud M, Billon-Grand G, Choquer M, et al. The Homeobox BcHOX8 Gene in Botrytis Cinerea Regulates Vegetative Growth and Morphology. PLoS ONE [Internet]. 2012 Oct 25 [cited 2018 Nov 29];7(10). Available from: https://www.ncbi.nlm.nih.gov/pmc/articles/PMC3485016/

17. Cary, Harris-Coward P, Scharfenstein L, Mack B, Chang PK, Wei Q, et al. The Aspergillus flavus Homeobox Gene, hbx1, Is Required for Development and Aflatoxin Production. Toxins [Internet]. 2017 Oct 12;9(10). Available from: http://www.mdpi.com/2072-6651/9/10/315

18. Arnaise S, Zickler D, Poisier C, Debuchy R. pah1: a homeobox gene involved in hyphal morphology and microconidiogenesis in the filamentous ascomycete Podospora anserina. Mol Microbiol. 2001 Jan;39(1):54–64.

19. Coppin E, Berteaux-Lecellier V, Bidard F, Brun S, Ruprich-Robert G, Espagne E, et al. Systematic deletion of homeobox genes in Podospora anserina uncovers their roles in shaping the fruiting body. PloS One. 2012;7(5):e37488.

20. Kim S, Park SY, Kim KS, Rho HS, Chi MH, Choi J, et al. Homeobox Transcription Factors Are Required for Conidiation and Appressorium Development in the Rice Blast Fungus Magnaporthe oryzae. Copenhaver GP, editor. PLoS Genet. 2009 Dec 4;5(12):e1000757.

21. Liu W, Xie S, Zhao X, Chen X, Zheng W, Lu G, et al. A homeobox gene is essential for conidiogenesis of the rice blast fungus Magnaporthe oryzae. Mol Plant-Microbe Interact MPMI. 2010 Apr;23(4):366–75.

22. Zheng W, Zhao X, Xie Q, Huang Q, Zhang C, Zhai H, et al. A Conserved Homeobox Transcription Factor Htf1 Is Required for Phialide Development and Conidiogenesis in Fusarium Species. Yu JH, editor. PLoS ONE. 2012 Sep 21;7(9):e45432.

23. Ghosh AK, Wangsanut T, Fonzi WA, Rolfes RJ. The GRF10 homeobox gene regulates filamentous growth in the human fungal pathogen Candida albicans. FEMS Yeast Res. 2015 Dec;15(8):fov093.

24. Calvo AM, Wilson RA, Bok JW, Keller NP. Relationship between Secondary Metabolism and Fungal Development. Microbiol Mol Biol Rev. 2002 Sep 1;66(3):447–59.

25. Calvo AM, Lohmar JM, Ibarra B, Satterlee T. 18 Velvet Regulation of Fungal Development. In: Wendland J, editor. Growth, Differentiation and Sexuality [Internet]. Cham: Springer International Publishing; 2016. p. 475–97. Available from: http://link.springer.com/10.1007/978-3-319-25844-7_18

26. Calvo AM, Cary JW. Association of fungal secondary metabolism and sclerotial biology. Front Microbiol. 2015;6:62.

27. Cary, Entwistle S, Satterlee T, Mack BM, Gilbert MK, Chang PK, et al. The Transcriptional Regulator Hbx1 Affects the Expression of Thousands of Genes in the Aflatoxin-Producing Fungus Aspergillus flavus. G3 Bethesda Md. 2019;9(1):167–78.

28. Satterlee T, Nepal B, Lorber S, Puel O, Calvo AM. The Transcriptional Regulator HbxA Governs Development, Secondary Metabolism, and Virulence in Aspergillus fumigatus. Appl Environ Microbiol. 2020 Jan 21;86(3). Available from: https://aem.asm.org/content/86/3/e01779-19

29. Calvo AM. The VeA regulatory system and its role in morphological and chemical development in fungi. Fungal Genet Biol. 2008 Jul;45(7):1053–61.

30. Cary, John E. Linz, Deepak Bhatnagar. Microbial Foodborne Diseases | Mechanisms of Pathogenesis and Toxin Sy. 2014. Available from: https://www.taylorfrancis.com/books/mono/10.1201/9781482278873/microbial-foodborne-diseases-jeffrey-cary-john-linz-deepak-bhatnagar

31. Han KH, Lee DB, Kim JH, Kim MS, Han KY, Kim WS, et al. Environmental factors affecting development of Aspergillus nidulans. J Microbiol. 2003;41(1):34–40.

32. Kosalková K, García-Estrada C, Ullán RV, Godio RP, Feltrer R, Teijeira F, et al. The global regulator LaeA controls penicillin biosynthesis, pigmentation and sporulation, but not roquefortine C synthesis in Penicillium chrysogenum. Biochimie. 2009 Feb;91(2):214–25.

33. Pandit, S. S., Satterlee, T., Nepal, B., Lorber, S., Puel, O., Espeso, E. A., et al. The transcriptional regulatory HbxA governs development and secondary metabolism in Aspergillus nidulans and Aspergillus fumigatus. 30th Fungal Genetics Conference, March 12-17, 2019, at Asilomar Conference Grounds in Pacific Grove, CA; 2019.

34. Son SH, Son YE, Cho HJ, Chen W, Lee MK, Kim LH, et al. Homeobox proteins are essential for fungal differentiation and secondary metabolism in Aspergillus nidulans. Sci Rep. 2020 Dec;10(1):6094.

35. Käfer E. Meiotic and Mitotic Recombination in Aspergillus and Its Chromosomal Aberrations. In: Caspari EW, editor. Advances in Genetics. Academic Press; 1977. p. 33–131. Available from: http://www.sciencedirect.com/science/article/pii/S006526600860245X

36. Feng X, Ramamoorthy V, Pandit SS, Prieto A, Espeso EA, Calvo AM. cpsA regulates mycotoxin production, morphogenesis and cell wall biosynthesis in the fungus Aspergillus nidulans. Mol Microbiol. 2017 Jul 1;105(1):1–24.

37. Ramamoorthy V, Dhingra S, Kincaid A, Shantappa S, Feng X, Calvo AM. The Putative C2H2 Transcription Factor MtfA Is a Novel Regulator of Secondary Metabolism and Morphogenesis in Aspergillus nidulans. PLOS ONE. 2013 Sep 16;8(9):e74122.

38. Szewczyk E, Nayak T, Oakley CE, Edgerton H, Xiong Y, Taheri-Talesh N, et al. Fusion PCR and gene targeting in *Aspergillus nidulans*. Nat Protoc. 2006 Dec;1(6):3111–20.

39. Stinnett SM, Espeso EA, Cobeño L, Araújo-Bazán L, Calvo AM. Aspergillus nidulans VeA subcellular localization is dependent on the importinlil? carrier and on light. Mol Microbiol. 2007 Jan;63(1):242– 55.

40. Krueger F. Trim Galore. 2022. Available from: https://github.com/FelixKrueger/TrimGalore

41. Wood DE, Lu J, Langmead B. Improved metagenomic analysis with Kraken 2. Genome Biol. 2019 Nov 28;20(1):257.

42. Basenko EY, Pulman JA, Shanmugasundram A, Harb OS, Crouch K, Starns D, et al. FungiDB: An Integrated Bioinformatic Resource for Fungi and Oomycetes. J Fungi Basel Switz. 2018 Mar 20;4(1):39.

43. Bushnell B, Rood J, Singer E. BBMerge - Accurate paired shotgun read merging via overlap. PloS One. 2017;12(10):e0185056.

44. Kim D, Paggi JM, Park C, Bennett C, Salzberg SL. Graph-based genome alignment and genotyping with HISAT2 and HISAT-genotype. Nat Biotechnol. 2019 Aug;37(8):907–15.

45. Li H, Handsaker B, Wysoker A, Fennell T, Ruan J, Homer N, et al. The Sequence Alignment/Map format and SAMtools. Bioinforma Oxf Engl. 2009 Aug 15;25(16):2078–9.

46. Pertea M, Pertea GM, Antonescu CM, Chang TC, Mendell JT, Salzberg SL. StringTie enables improved reconstruction of a transcriptome from RNA-seq reads. Nat Biotechnol. 2015 Mar;33(3):290–5.

47. Love MI, Huber W, Anders S. Moderated estimation of fold change and dispersion for RNA-seq data with DESeq2. Genome Biol. 2014;15(12):550.

48. Priebe S, Kreisel C, Horn F, Guthke R, Linde J. FungiFun2: a comprehensive online resource for systematic analysis of gene lists from fungal species. Bioinforma Oxf Engl. 2015 Feb 1;31(3):445–6.

49. Lind AL, Wisecaver JH, Smith TD, Feng X, Calvo AM, Rokas A. Examining the Evolution of the Regulatory Circuit Controlling Secondary Metabolism and Development in the Fungal Genus Aspergillus. PLOS Genet. 2015 Mar 18;11(3):e1005096.

50. Inglis DO, Binkley J, Skrzypek MS, Arnaud MB, Cerqueira GC, Shah P, et al. Comprehensive annotation of secondary metabolite biosynthetic genes and gene clusters of Aspergillus nidulans, A. fumigatus, A. niger and A. oryzae. BMC Microbiol. 2013 Apr 26;13(1):91.

51. Etxebeste O. Transcription Factors in the Fungus Aspergillus nidulans: Markers of Genetic Innovation, Network Rewiring and Conflict between Genomics and Transcriptomics. J Fungi. 2021 Jul 25;7(8):600.

52. Cleveland DW. Peptide mapping in one dimension by limited proteolysis of sodium dodecyl sulfate-solubilized proteins. In: Methods in Enzymology. Academic Press; 1983. p. 222–9. (Biomembranes Part J: Membrane Biogenesis: Assembly and Targeting (General Methods, Eukaryotes); vol. 96). Available from: https://www.sciencedirect.com/science/article/pii/S0076687983960202

53. Robert X, Gouet P. Deciphering key features in protein structures with the new ENDscript server. Nucleic Acids Res. 2014 Jul 1;42(W1):W320–4.

54. Arnaud MB, Cerqueira GC, Inglis DO, Skrzypek MS, Binkley J, Chibucos MC, et al. The Aspergillus Genome Database (AspGD): recent developments in comprehensive multispecies curation, comparative genomics and community resources. Nucleic Acids Res. 2012 Jan;40(Database issue):D653–9.

55. Adams TH, Boylan MT, Timberlake WE. brlA is necessary and sufficient to direct conidiophore development in aspergillus nidulans. Cell. 1988 Jul 29;54(3):353–62.

56. Andrianopoulos A, Timberlake WE. The Aspergillus nidulans abaA gene encodes a transcriptional activator that acts as a genetic switch to control development. Mol Cell Biol. 1994 Apr;14(4):2503– 15.

57. Chang YC, Timberlake WE. Identification of Aspergillus brlA response elements (BREs) by genetic selection in yeast. Genetics. 1993 Jan 1;133(1):29–38.

58. Prade RA, Timberlake WE. The Aspergillus nidulans brlA regulatory locus consists of overlapping transcription units that are individually required for conidiophore development. EMBO J. 1993 Jun;12(6):2439–47.

59. Etxebeste O, Ni M, Garzia A, Kwon NJ, Fischer R, Yu JH, et al. Basic-Zipper-Type Transcription Factor FlbB Controls Asexual Development in Aspergillus nidulans. Eukaryot Cell. 2008 Jan;7(1):38–48.

60. Son YE, Cho HJ, Lee MK, Park HS. Characterizing the role of Zn cluster family transcription factor ZcfA in governing development in two Aspergillus species. PLoS ONE. 2020 Feb 4;15(2):e0228643.

61. Smith TD, Calvo AM. The mtfA transcription factor gene controls morphogenesis, gliotoxin production, and virulence in the opportunistic human pathogen Aspergillus fumigatus. Eukaryot Cell. 2014 Jun;13(6):766–75.

62. Zhuang Z, Lohmar JM, Satterlee T, Cary JW, Calvo AM. The Master Transcription Factor mtfA Governs Aflatoxin Production, Morphological Development and Pathogenicity in the Fungus Aspergillus flavus. Toxins. 2016 Jan 20;8(1):29.

63. Pandit SS, Lohmar JM, Ahmed S, Etxebeste O, Espeso EA, Calvo AM. UrdA Controls Secondary Metabolite Production and the Balance between Asexual and Sexual Development in Aspergillus nidulans. Genes. 2018 Nov 23;9(12). Available from: https://www.ncbi.nlm.nih.gov/pmc/articles/PMC6316066/

64. Thieme KG, Gerke J, Sasse C, Valerius O, Thieme S, Karimi R, et al. Velvet domain protein VosA represses the zinc cluster transcription factor SclB regulatory network for Aspergillus nidulans asexual development, oxidative stress response and secondary metabolism. PLOS Genet. 2018 Jul 25;14(7):e1007511.

65. Alkahyyat F, Ni M, Kim SC, Yu JH. The WOPR Domain Protein OsaA Orchestrates Development in Aspergillus nidulans. PLOS ONE. 2015 Sep 11;10(9):e0137554.

66. Bayram O, Krappmann S, Ni M, Bok JW, Helmstaedt K, Valerius O, et al. VelB/VeA/LaeA complex coordinates light signal with fungal development and secondary metabolism. Science. 2008 Jun 13;320(5882):1504–6.

67. Arst HN, Hondmann DHA, Visser J. A translocation activating the cryptic nitrogen regulation gene areB inactivates a previously unidentified gene involved in glycerol utilisation in Aspergillus nidulans | SpringerLink. 1990. Available from: https://link.springer.com/article/10.1007/BF00315805

68. Kowalczyk JE, Gruben BS, Battaglia E, Wiebenga A, Majoor E, de Vries RP. Genetic Interaction of Aspergillus nidulans galR, xlnR and araR in Regulating D-Galactose and L-Arabinose Release and Catabolism Gene Expression. PLoS ONE. 2015 Nov 18;10(11):e0143200.

69. Strauss J, Horvath HK, Abdallah BM, Kindermann J, Mach RL, Kubicek CP. The function of CreA, the carbon catabolite repressor of Aspergillus nidulans, is regulated at the transcriptional and post-transcriptional level. Mol Microbiol. 1999 Apr;32(1):169–78.

70. Askenazi M, Driggers EM, Holtzman DA, Norman TC, Iverson S, Zimmer DP, et al. Integrating transcriptional and metabolite profiles to direct the engineering of lovastatin-producing fungal strains. Nat Biotechnol. 2003 Feb;21(2):150–6.

71. Ukil L, Varadaraj A, Govindaraghavan M, Liu HL, Osmani SA. Copy Number Suppressors of the Aspergillus nidulans nimA1 Mitotic Kinase Display Distinctive and Highly Dynamic Cell Cycle-Regulated Locations. Eukaryot Cell. 2008 Dec;7(12):2087–99.

72. Dyer PS, O’Gorman CM. Sexual development and cryptic sexuality in fungi: insights from Aspergillus species. FEMS Microbiol Rev. 2012 Jan 1;36(1):165–92.

73. Pyrzak W, Miller KY, Miller BL. Mating Type Protein Mat1-2 from Asexual Aspergillus fumigatus Drives Sexual Reproduction in Fertile Aspergillus nidulans. Eukaryot Cell. 2008 Jun;7(6):1029–40.

74. Piłsyk S, Natorff R, Sieńko M, Skoneczny M, Paszewski A, Brzywczy J. The Aspergillus nidulans metZ gene encodes a transcription factor involved in regulation of sulfur metabolism in this fungus and other Eurotiales. Curr Genet. 2015 May;61(2):115–25.

75. Xiong Y, Wu VW, Lubbe A, Qin L, Deng S, Kennedy M, et al. A fungal transcription factor essential for starch degradation affects integration of carbon and nitrogen metabolism. PLOS Genet. 2017 May 3;13(5):e1006737.

76. Ahmed YL, Gerke J, Park HS, Bayram Ö, Neumann P, Ni M, et al. The Velvet Family of Fungal Regulators Contains a DNA-Binding Domain Structurally Similar to NF-κB. Stock AM, editor. PLoS Biol. 2013 Dec 31;11(12):e1001750.

77. Röhrig J, Yu Z, Chae KS, Kim JH, Han KH, Fischer R. The Aspergillus nidulans Velvet-interacting protein, VipA, is involved in light-stimulated heme biosynthesis. Mol Microbiol. 2017;105(6):825–38.

78. Alaminos, Ramos. The methionine biosynthetic pathway from homoserine in Pseudomonas putida involves the metW, metX, metZ, metH and metE gene products - PubMed. 2001. Available from: https://pubmed.ncbi.nlm.nih.gov/11479715/

79. Chiang YM, Szewczyk E, Davidson AD, Entwistle R, Keller NP, Wang CCC, et al. Characterization of the Aspergillus nidulans Monodictyphenone Gene Cluster. Appl Env Microbiol. 2010 Apr 1;76(7):2067–74.

80. Andersen MR, Nielsen JB, Klitgaard A, Petersen LM, Zachariasen M, Hansen TJ, et al. Accurate prediction of secondary metabolite gene clusters in filamentous fungi. Proc Natl Acad Sci U S A. 2013 Jan 2;110(1):E99–107.

81. Lo HC, Entwistle R, Guo CJ, Ahuja M, Szewczyk E, Hung JH, et al. Two separate gene clusters encode the biosynthetic pathway for the meroterpenoids austinol and dehydroaustinol in Aspergillus nidulans. J Am Chem Soc. 2012 Mar 14;134(10):4709–20.

82. Rokas A, Mead ME, Steenwyk JL, Raja HA, Oberlies NH. Biosynthetic gene clusters and the evolution of fungal chemodiversity. Nat Prod Rep. 2020 Jul 1;37(7):868–78.

83. Chanda A, Roze LV, Kang S, Artymovich KA, Hicks GR, Raikhel NV, et al. A key role for vesicles in fungal secondary metabolism. Proc Natl Acad Sci. 2009 Nov 17;106(46):19533–8.

84. Ali, Marco I Ries, Peter P Lankhorst, Rob A M van der Hoeven, Olaf L Schouten, Marek Noga, et al. A non-canonical NRPS is involved in the synthesis of fungisporin and related hydrophobic cyclic tetrapeptides in Penicillium chrysogenum - PubMed. 2014. Available from: https://pubmed.ncbi.nlm.nih.gov/24887561/

85. Himaja, M., Elizabeth, L., Moonjit, D., Asif, K. Synthesis and antimicrobial evaluation of N-methylated analog of fungisporin. Univ J Pharm. 2015;4:10–4.

86. Gozari M, Alborz M, El-Seedi HR, Jassbi AR. Chemistry, biosynthesis and biological activity of terpenoids and meroterpenoids in bacteria and fungi isolated from different marine habitats. Eur J Med Chem. 2021 Jan 15;210:112957.

87. Fuloria NK, Raheja RK, Shah KH, Oza, M.J. Biological activities of meroterpenoids isolated from different sources. Frontiers Media SA; 2022. 174 p.

88. Matsuda Y, Awakawa T, Mori T, Abe I. Unusual chemistries in fungal meroterpenoid biosynthesis. Curr Opin Chem Biol. 2016 Apr 1;31:1–7.

